# A DNA damage-activated kinase controls bacterial immune pathway expression

**DOI:** 10.64898/2026.02.02.703251

**Authors:** Lydia R. Chambers, Phoolwanti Rani, Ryan K Min, Elizabeth Villa, Kevin D. Corbett

## Abstract

Bacteria encode myriad stress-response pathways that protect their hosts against both internal and external threats. A key question is how these pathways are regulated, especially anti-phage immune pathways that mediate host cell killing. Here, we identify two proteins termed CapK and CapS that are encoded upstream of diverse immune operons, and regulate their expression in response to DNA damage. CapK resembles bacterial anti-sigma factor kinases, and CapS resembles these proteins’ STAS domain antagonists. CapS is a DNA-binding transcriptional repressor, and phosphorylation of CapS by CapK results in dissociation of a CapS homodimer and de-repression of transcription. CapK’s kinase activity is directly activated by single-stranded DNA generated as a by-product of DNA repair. Finally, we show that CapK and CapS-like proteins have been co-opted into an anti-phage toxin-antitoxin system with a VapC-like protein, where they similarly respond to DNA damage to activate VapC’s nuclease activity. Overall, our results reveal how a kinase-substrate pair can regulate expression of an adjacent operon in response to DNA damage, and highlight the modularity of immune and other stress-response pathways.

## INTRODUCTION

Under the pressure of environmental stress and the constant threat of phage infection, bacteria have evolved a large repertoire of immune pathways with diverse mechanisms for infection sensing and host protection (Bernheim & Sorek, 2020; Hampton *et al*, 2020). In the past decade, hundreds of distinct bacterial immune pathways have been identified, which are often encoded in clusters termed “defense islands” in bacterial genomes. Immune pathways typically function either by directly targeting foreign DNA (as in Restriction-Modification (R-M) and CRISPR-Cas systems), or by causing growth inhibition and/or cell death to limit phage replication (broadly termed abortive infection) (Gao *et al*, 2020; Johnson *et al*, 2022; Lopatina *et al*, 2020; Tamulaitiene *et al*, 2024). Abortive infection pathways in particular must be tightly regulated to prevent aberrant activation, which could lead to toxicity in the absence of infection.

Many antiphage immune pathways are regulated at the level of transcription and/or translation, with expression of these pathways induced by external or internal stress signals. For example, quorum sensing is utilized by some CRISPR-Cas systems, with these pathways’ expression repressed at low cell density and de-repressed at high cell density (Høyland-Kroghsbo *et al*, 2017; Patterson *et al*, 2016). Many R-M systems encode a controller (C) protein that enables fine-tuned control over the relative expression timing of these systems’ methylase and endonuclease components (Negri *et al*, 2021). Additionally, two families of DNA damage-responsive transcriptional regulators have been identified that regulate the expression of adjacently-encoded immune operons, including BREX, Pycsar, CBASS, and DISARM. These include CapW/BrxR, which represses transcription until it binds single-stranded DNA (ssDNA); and CapH+CapP, whose CapP protease is activated upon ssDNA binding and cleaves the DNA-binding repressor CapH (Lau *et al*, 2022; Blankenchip *et al*, 2022; Blankenchip & Corbett, 2024; Picton *et al*, 2022; Luyten *et al*, 2022). These two families of DNA-damage activated regulators highlight the importance of direct control of gene expression in immune pathways.

Here, we identify and characterize a third family of immune pathway-associated, DNA-damage activated transcriptional regulators termed CapK+CapS. We identify *capK+capS* genes associated with diverse known and putative bacterial immune pathways, including CBASS, Bil (bacterial ISG15-like), and Bub (bacterial ubiquitination-like). We find that CapS is a DNA-binding transcriptional repressor, which is phosphorylated by CapK upon DNA damage to de-repress an adjacently-encoded immune operon. Finally, we identify standalone operons encoding CapK and CapS-like proteins plus a VapC-like RNase toxin, and show that these operons control VapC activation through a similar DNA damage sensing mechanism. Overall, our findings emphasize the prevalence and importance of immune pathway regulation at the transcriptional level, and the central role of DNA damage in activation of immune pathway expression.

## RESULTS

### A predicted kinase-substrate gene pair associated with bacterial immune operons

We recently defined a family of putative bacterial immune pathways termed Bub (<u>b</u>acterial <u>ub</u>iquitination-like), and identified hundreds of Bub operons in diverse bacteria (Gong *et al*, 2025; Ye *et al*, 2025). By manually inspecting these operons’ gene neighborhoods, we found that 219 out of 544 identified Bub operons are associated with either CapW (<u>C</u>BASS-<u>a</u>ssociated <u>p</u>rotein, <u>W</u>YL domain) or CapH+CapP (<u>C</u>BASS-<u>a</u>ssociated <u>p</u>roteins, <u>H</u>elix-turn-helix and <u>P</u>eptidase), both of which control the expression of their adjacent operon in response to DNA damage (Blankenchip *et al*, 2022; Blankenchip & Corbett, 2024; Lau *et al*, 2022). Further inspection revealed that 59 of 544 Bub operons were associated with two genes of unknown function, one of which encodes predicted HTH (<u>h</u>elix-turn <u>h</u>elix), STAS (<u>s</u>ulphate transporter and <u>a</u>nti-<u>s</u>igma factor antagonist) (Aravind & Koonin, 2000), and GHKL (DNA <u>G</u>yrase, <u>h</u>istidine <u>k</u>inase, Mut<u>L</u>) (Dutta & Inouye, 2000) ATPase/kinase domains, and the second of which encodes a predicted STAS domain followed by a wHTH (<u>w</u>inged <u>h</u>elix-turn-<u>h</u>elix) domain (**Figure 1A, Table S1**). These two genes’ location adjacent to Bub operons’ promoters suggests that they may regulate transcription in a manner similar to CapW and CapH+CapP. Following a similar naming convention, we named the HTH-STAS-GHKL kinase protein CapK (<u>K</u>inase), and the STAS-wHTH protein CapS (<u>S</u>TAS domain). Reflecting the fact that CapK+CapS was identified adjacent to Bub immune pathways rather than CBASS, we propose that “Cap” for transcriptional regulators (including CapW, CapH+CapP, and CapK+CapS) henceforth stand for “<u>C</u>ontroller of <u>a</u>ssociated/adjacent <u>p</u>athways”.

**Figure 1.**
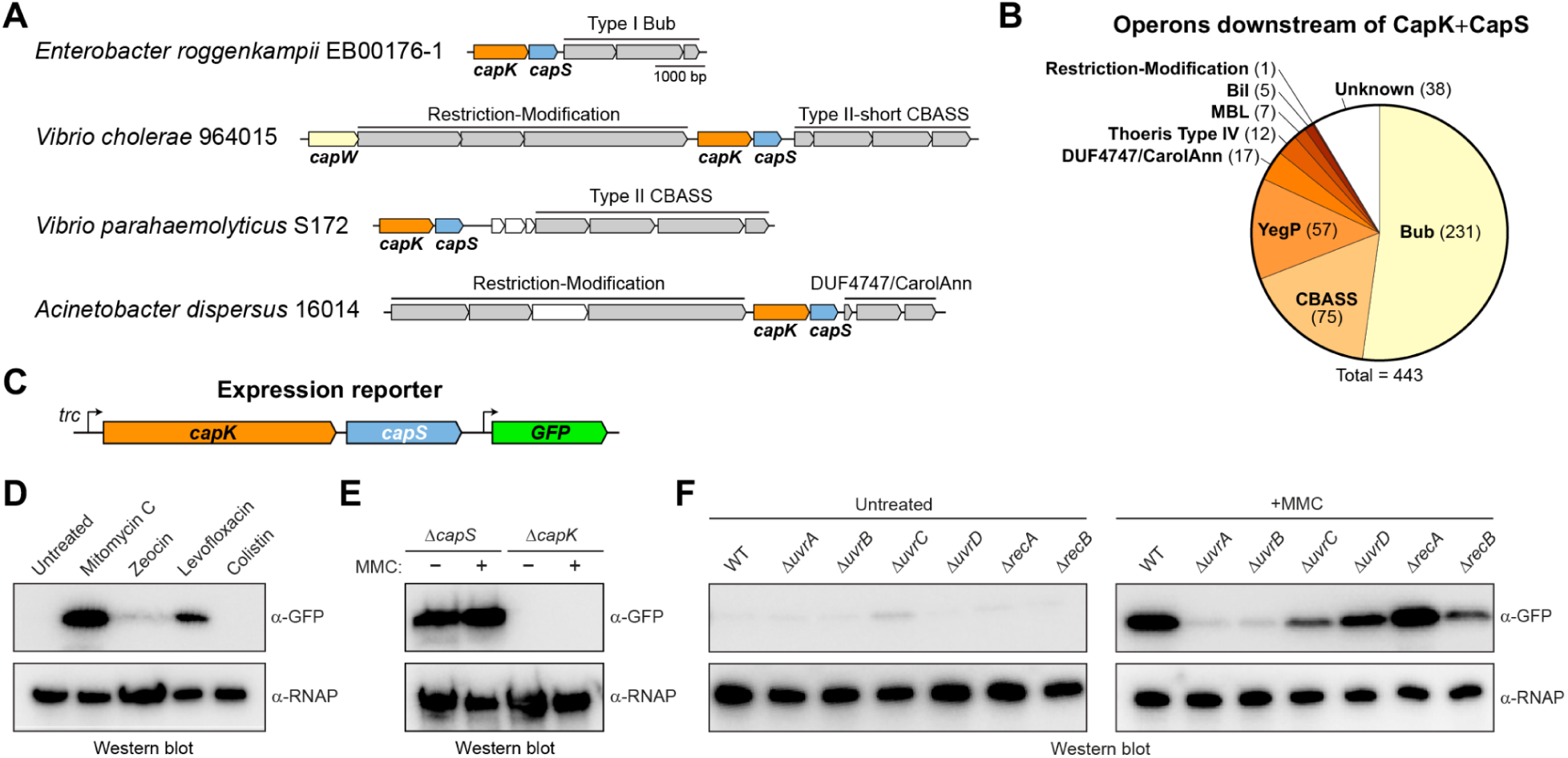
Identification of CapK and CapS. (A) Operon schematics of four bacterial immune pathways (pink) associated with *capK* (orange) and *capS* (blue) genes. Immune pathways are shown in gray and labeled; unknown genes are shown in white. See **Table S2** for full list and accession numbers. (B) Classification of 443 identified *capK*/*capS*-associated operons. (C) Schematic of the GFP expression reporter derived from an *E. roggenkampii* strain EB00176-1 CBASS-associated *capKS* locus (see panel A). An IPTG-inducible *trc* promoter is positioned upstream of *capK*, and the CBASS operon is replaced by the coding sequence for GFP (green). For some experiments (noted in figure legends), an N-terminal FLAG tag was fused to the *capS* gene. (D) Anti-GFP western blot showing induction of GFP expression in the presence of different small molecules (concentrations noted in **Materials and Methods**), using the GFP expression reporter expressing FLAG-CapS. α-RNAP: loading control blot for RNA polymerase subunit RpoS. (E) Anti-GFP western blot showing the effect of deleting *capS* or *capK* in the GFP expression reporter, in the presence or absence of mitomycin C (MMC). (F) Anti-GFP western blot showing GFP expression in the presence or absence of mitomycin C (MMC) in the indicated single-gene knockout *E. coli* strains. This experiment used the GFP expression reporter expressing FLAG-CapS.

We performed comprehensive BLAST searches in the Integrated Microbial Genomes (IMG) database, and identified 443 homologs of *capK*, all of which are immediately followed by *capS*. In 346 of 443 cases, *capK* and *capS* are followed by a known or predicted bacterial immune operon (**Figure 1B**, **Table S2**), including CBASS (72 cases) (Davies *et al*, 2012; Cohen *et al*, 2019; Ye *et al*, 2020), DUF4747/CarolAnn prophage system (17 cases) (Montgomery *et al*, 2019; Parma *et al*, 1992), and Type IV Thoeris (12 cases) (Rousset *et al*, 2025). We also identified five Bil operons (Millman *et al*, 2022; Chambers *et al*, 2024; Hör *et al*, 2024), 7 MBL operons related to Type II CBASS, and 230 Bub operons with upstream *capK* and *capS*. In 52 cases, *capK* and *capS* were followed by a gene encoding a protein homologous to the uncharacterized *E. coli* protein YegP. Finally, we found that in 88 out of 443 cases, *capK* and *capS* are located immediately downstream of a predicted R-M system (**Figure 1A**), suggesting that the two genes may coordinate the immune activities of an upstream R-M system and a downstream immune system like CBASS or Bub.

### CapK and CapS regulate transcription of a downstream immune operon

The location of *capK* and *capS* upstream of diverse immune operons suggests that, like CapW and CapH+CapP, they regulate transcription of the downstream operon in response to stress. To test this idea, we chose a *capK*+*capS* gene pair associated with a Bub operon in *Enterobacter roggenkampii* strain EB00176-1 (**Figure 1A**), and created a GFP expression reporter by replacing the Bub operon with a gene encoding GFP (maintaining the intergenic sequence downstream of *capS*; **Figure 1C**). We also fused *capS* to an N-terminal FLAG epitope tag, enabling us to track expression of CapS directly (see below). In log-phage *E. coli* cultures in rich media, GFP expression was undetectable. When we exposed cells to different types of stress by adding antibiotics, membrane-disrupting compounds, and DNA-damaging agents, we observed induction of GFP expression after adding the DNA damaging agent mitomycin C (MMC) or the DNA gyrase inhibitor levofloxacin (**Figure 1D**). We next deleted either *capK* or *capS* in our GFP expression reporter, and found that deletion of *capS* results in constitutive GFP expression, and that deletion of *capK* disrupts MMC-activated expression (**Figure 1E**). These data suggest that CapS is a transcriptional repressor for the downstream operon, and that CapK relieves CapS-mediated repression in response to DNA damage.

Mitomycin C damages DNA by creating interstrand crosslinks (Tomasz, 1994). These lesions are predominantly repaired by the UvrABCD nucleotide excision repair (NER) pathway (Truglio *et al*, 2006). To determine whether the UvrABCD pathway plays a role in CapK+CapS mediated transcriptional regulation, we tested MMC-dependent GFP reporter expression in *E. coli* strains lacking *uvrA*, *uvrB*, *uvrC*, or *uvrD*, in addition to strains lacking the homologous recombination repair proteins *recA* or *recB*. We found that MMC-dependent GFP expression depends on both *uvrA* and *uvrB*, which act early in the NER pathway to recognize damaged DNA (**Figure 1F**). These data support a model in which CapK and CapS sense a byproduct of NER to activate downstream operon expression.

### CapS binds two sites in the *capK-capS* region

Our GFP expression reporter assays suggested that CapS acts as a repressor, likely by binding promoter DNA through its predicted wHTH domain. To identify CapS binding sites, we modified our GFP expression reporter to include the entire *E. roggenkampii capK-capS* region, including 660 bp upstream of *capK* and the entire intergenic region between *capS* and the downstream Bub operon. We used ChIP-Seq to measure binding of FLAG-CapS to DNA, in either unperturbed cells or one hour after MMC treatment (**Figure 2A**). We detected two strong peaks of CapS-DNA binding: one in the short intergenic region between *capK* and *capS* (peak #1) and a second in the longer intergenic region between *capS* and the downstream GFP gene (peak #2). Exposure of cells to MMC did not result in a measurable reduction of CapS-DNA binding (**Figure 2A**), suggesting either that CapS is not fully removed from DNA upon exposure of cells to MMC, or that CapS-DNA binding recovers by one hour after MMC treatment.

**Figure 2.**
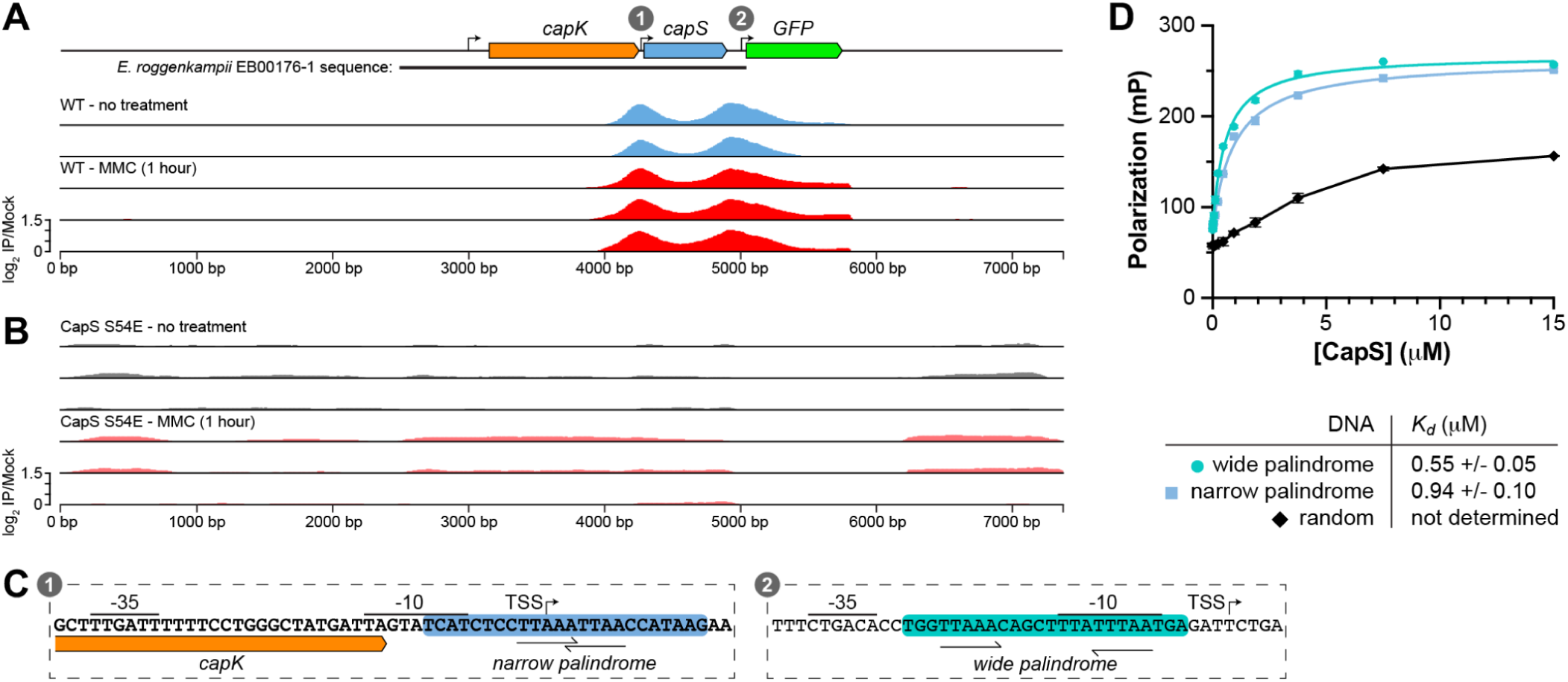
CapS binds two sites in the *capK-capS* region. (A) ChIP-Seq results measuring binding of FLAG-CapS to a plasmid encoding the *capK*-*capS* genomic region (660 bp upstream of *capK* to the start codon of the downstream Bub operon) followed by a gene coding for GFP. Data is expressed as the log2 enrichment of FLAG-immunopurified samples over mock-purified samples (log2 IP/Mock), normalized for RPM coverage (see **Methods**). No-treatment samples are shown in blue, and MMC-treated samples are shown in red. (B) ChIP-Seq results measuring binding of FLAG-CapS S54E to a plasmid encoding the *capK*-*capS* genomic region followed by a gene coding for GFP. No-treatment samples are shown in gray, and MMC-treated samples are shown in pink. (C) Closeup schematics of promoters #1 (upstream of *capS*) and #2 (downstream of *capS*) as indicated in panel (A). (D) Fluorescence polarization DNA binding assay with *E. roggenkampii* CapS and three 24-bp dsDNAs: the “narrow palindrome” identified in promoter #1 (blue squares), the “wide palindrome” identified in promoter #2 (green circles), and a random sequence (black diamonds). Each datapoint is an average of three technical replicates, and error bars indicate the mean +/- standard deviation (error bars not shown if they are smaller than the datapoint itself). Fit *K_d_* values are shown in the table at bottom.

We used the BPROM server (Solovyev & Salamov, 2011) to identify putative promoter sequences in both the *capK-capS* intergenic region and the *capS*-downstream gene intergenic region (**Figure 2C**). Based on our observation that purified *E. roggenkampii* CapS is dimeric in solution (**Figure S1**), we next searched for palindromic sequences overlapping these promoters and corresponding to ChIP-Seq peaks #1 and #2 (**Figure 2C**). We identified palindromic sequences in both putative promoters with the same half-site sequence (TTAAA), but with the half-sites spaced differently in the two promoters. In the *capK-capS* intergenic region (corresponding to peak #1), we identified a “narrow palindrome” with the sequence TTAAATTAA, where the two half-sites overlap by one base pair (the reverse half-site reads TTAAT in this case). In the *capS*-downstream gene region (corresponding to peak #2), we identified a “wide palindrome” with the sequence TTAAAcagctttaTTTAA, where the two half-sites are separated by eight base pairs. We performed fluorescence polarization DNA binding assays with 24-bp DNA segments containing both palindromic sequences, and found that CapS binds the wide palindrome with a dissociation constant (*K_d_*) of 0.55 +/- 0.05 µM (**Figure 2D**), and binds the narrow palindrome with a slightly lower affinity (*K_d_*= 0.95 +/- 0.10 µM; **Figure 2D**). Demonstrating specificity, CapS did not detectably bind a random-sequence DNA of the same length (**Figure 2D**). Overall, these data show that CapS can bind palindromic sequences with a half-site sequence of TTAAA, with these half-sites spaced either eight base pairs apart or overlapping by one base pair. Combined with our ChIP-Seq data, these binding data further suggest that CapS regulates both its own expression and that of the downstream operon.

### The structure of CapS reveals two distinct dimer interfaces

We next crystallized and determined the structure of *E. roggenkampii* CapS to a resolution of 1.55 Å (**Table S5**). The structure reveals that CapS comprises an N-terminal STAS domain and a C-terminal wHTH domain (**Figure 3A-B**). Structure comparisons with the DALI server show strong similarity of CapS’s wHTH domain to DNA-binding transcriptional repressors, including ArsR (Viswanathan *et al*, 2021; Prabaharan *et al*, 2019) and MarR-family proteins (Conway *et al*, 2022; Peng *et al*, 2017; Song *et al*, 2024) (**Figure S2A-B**). We modeled CapS bound to DNA based on a structure of a DNA-bound MarR transcription factor (Song *et al*, 2024), and identified several absolutely-conserved residues predicted to contact DNA, including N145, K151, and two glycine residues (169-170) that are positioned at the distal end of the wing motif predicted to bind the DNA minor groove (**Figure 3C, S2B**). Mutation of N145 to alanine (N145A) eliminated binding to the wide palindrome DNA, supporting a model in which CapS binds DNA via its wHTH domain (**Figure 3D**).

**Figure 3.**
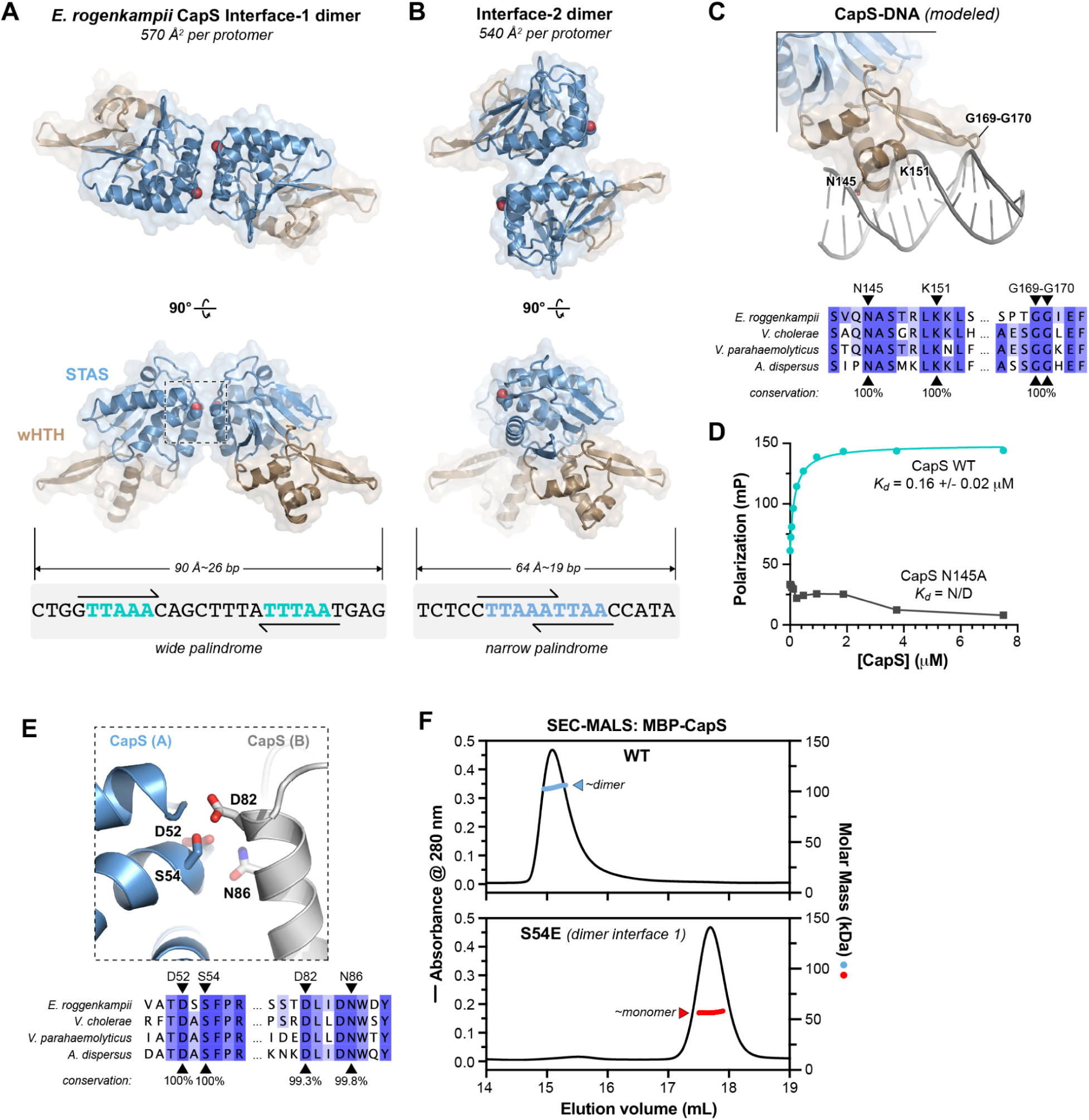
Structure of *E. roggenkampii* CapS. (A) *Top:* Top and side views of the *E. roggenkampii* CapS interface-1 dimer, with N-terminal STAS domain (residues 1-111) blue and C-terminal wHTH domain (residues 112-184) brown. Residue Ser54, buried in the dimer interface, is shown as spheres. *Bottom:* “wide palindrome” sequence from promoter #2 (Figure 2C), to scale with CapS. (B) *Top:* Top and side views of the *E. roggenkampii* CapS interface-2 dimer, with N-terminal STAS domain (residues 1-111) blue and C-terminal wHTH domain (residues 112-184) brown. *Bottom:* “narrow palindrome” sequence from promoter #1 (Figure 2C), to scale with CapS. (C) *Top:* Model of the CapS wHTH domain (brown) bound to DNA (gray), generated by overlaying the CapS wHTH domain with a DNA-bound structure of MarR (PDB ID 8YLG; **Figure S2B**) (Song *et al*, 2024). Conserved residues predicted to bind DNA (N145, K151, G169, G170) are shown as sticks and labeled. *Bottom:* Section of sequence alignment the four CapS proteins shown in Figure 1A (*E. roggenkampii* (NCBI WP_001567865.1), *V. cholerae* (NCBI WP_046127252.1), *V. parahaemolyticus* (IMG 2662856252), showing conservation of predicted DNA-binding residues. (D) Fluorescence polarization DNA binding assay with *E. roggenkampii* CapS (wild-type shown in blue circles, N145A shown in dark gray squares) and the “wide palindrome” from promoter #2. Each datapoint is an average of three technical replicates, and arrow bars indicate the mean +/- standard deviation (error bars not shown if they are smaller than the datapoint itself). (E) *Top:* Closeup of CapS residue Ser54, which is buried in the CapS interface-1 dimer and is surrounded by several negatively-charged or polar residues from the opposite CapS protomer (gray). *Bottom:* Section of sequence alignment the four CapS proteins shown in Figure 1A (*E. roggenkampii* (NCBI WP_001567865.1), *V. cholerae* (NCBI WP_046127252.1), *V. parahaemolyticus* (IMG 2662856252), showing conservation of residues in dimer interface 1. (F) SEC-MALS of purified *E. roggenkampii* MBP-tagged CapS wild-type (top) or S54E (bottom). CapS wild-type shows a molecular weight consistent with a homodimer (128.4 kDa), and the S54E mutant shows a molecular weight consistent with a monomer (64.2 kDa).

Our structure of *E. roggenkampii* CapS contains one protomer in the crystallographic asymmetric unit, but the protein exists as a homodimer in solution (**Figure S1**) and binds palindromic DNA sequences. We used the PDBePISA server (Krissinel & Henrick, 2007) to identify crystal packing interactions likely to represent the physiologically relevant dimer interface. We identified two major crystal packing interfaces, both involving the STAS domain: interface 1 buries 570 Å^2^ of surface area per protomer (**Figure 3A**), and interface 2 buries 540 Å^2^ of surface area per protomer (**Figure 3B**). Revisiting the two identified binding sites for *E. roggenkampii* CapS, we found that the spacing of the wide palindrome perfectly matches the spacing of wHTH domains in a CapS dimer generated by interface 1 (**Figure 3A***, bottom*). Similarly, the spacing of the narrow palindrome perfectly matches the spacing of a CapS dimer generated by interface 2 (**Figure 3B***, bottom*).

We next crystallized and determined two independent structures of *V. cholerae* CapS to a resolution of 2.38 Å (form 1; space group P3_1_21) and 1.84 Å (form 2; space group C222_1_), respectively. *V. cholerae* CapS is 41% identical to *E. roggenkampii* CapS, and the two proteins overlay closely with an overall Cα r.m.s.d. of 2.3 Å (over 176 Cα pairs; **Figure S3A**). The two structures of *V. cholerae* CapS each contain five CapS protomers per asymmetric unit, and both structures show crystal packing interactions equivalent to both interface 1 and interface 2 in our structure of *E. roggenkampii* CapS (**Figure S3B-C**). Moreover, since dimer interfaces 1 and 2 do not overlap, a single CapS protomer can associate with two other CapS protomers via these two interfaces. In this manner, a continuous helical filament of CapS could self-assemble using these interfaces, with all protomers’ wHTH domains aligned to bind a single continuous DNA duplex (**Figure S3D**).

### CapK and CapS resemble bacterial anti-sigma factors and their antagonists

Structure comparisons with the DALI server show that the N-terminal STAS domain of CapS is most similar to bacterial anti-sigma factor antagonists (also known as anti-anti-sigma factors) from *Bacillus* (SpoIIAA) (Masuda *et al*, 2004) and Mycobacteria (RsfB; PDB ID 8IH8 *unpublished*) (**Figure S2C-D**), which regulate important developmental switches in processes like sporulation and stress response (Dworkin & Losick, 2001; Moy & Seshu, 2021). These proteins contain a single STAS domain with a conserved serine residue that is phosphorylated by a GHKL-family histidine kinase domain in their cognate anti-sigma factors (*Bacillus* SpoIIAB and *Mycobacteria* RsbW, respectively). We found that CapS possesses an absolutely conserved serine in an equivalent position on the second α-helix of its STAS domain (*E. roggenkampii* CapS S54; **Figure 3E**). To test whether phosphorylation affects the oligomerization state of CapS in solution, we generated an *E. roggenkampii* CapS mutant with S54 mutated to glutamate (S54E), mimicking phosphorylation of this residue. By size exclusion chromatography coupled to multi-angle light scattering (SEC-MALS), we found that while wild-type CapS is dimeric, CapS S54E is monomeric (**Figure 3F**). In our structure of *E. roggenkampii* CapS, S54 is buried in dimer interface 1, and surrounded by highly conserved polar/negatively-charged residues (**Figure 3A,E**), whereas this residue is not involved in dimer interface 2. The fact that the S54E mutant disrupts the CapS dimer observed in solution therefore indicates that CapS dimerizes primarily through interface 1.

To assess how phosphorylation of CapS S54 affects DNA binding, we performed ChIP-Seq analysis. Strikingly, we found that mutating S54 to glutamate (mimicking phosphorylation) resulted in a complete loss of binding to both sites in the *capK-capS* region (**Figure 2B**). Overall, these data support a model for CapS-DNA binding in which (1) “wide palindrome” sites can be directly bound by a CapS interface-1 dimer; (2) “narrow palindrome” sites may recruit two CapS interface-1 dimers that each bind one half-site of the narrow palindrome, and that associate with the neighboring CapS dimer through interface 2. In both cases, further oligomerization could occur through recruitment of additional CapS dimers to adjacent sites along the DNA. Indeed, such oligomerization may explain why the CapS DNA binding peaks we observe by ChIP-Seq are notably wider than would be expected from the post-shearing DNA fragment length of ∼200 bp used in these experiments (**Figure 2A**).

Next, we sought to understand how CapK might bind and phosphorylate CapS. While we could purify a CapK-CapS complex with apparent 1:1 stoichiometry (**Figure S4A**), we could not determine an experimental structure of this complex. Therefore, we used AlphaFold 3 to predict the structure of *E. roggenkampii* CapK bound to ATP and CapS (**Figure 4A-B, Figure S4B-C**). The two proteins are strongly predicted to bind one another (AlphaFold 3 ipTM = 0.91; values above 0.8 represent highly-confident interaction predictions (Kim *et al*, 2024)). In the resulting model, CapS S54 is positioned adjacent to the CapK histidine kinase active site, poised for phosphorylation (**Figure 4B**). This predicted structure closely matches prior structures of *Bacillus* and Mycobacteria anti-sigma factor antagonists bound to their cognate kinase anti-sigma factors (**Figure 4C, S4D**) (Masuda *et al*, 2004).

**Figure 4.**
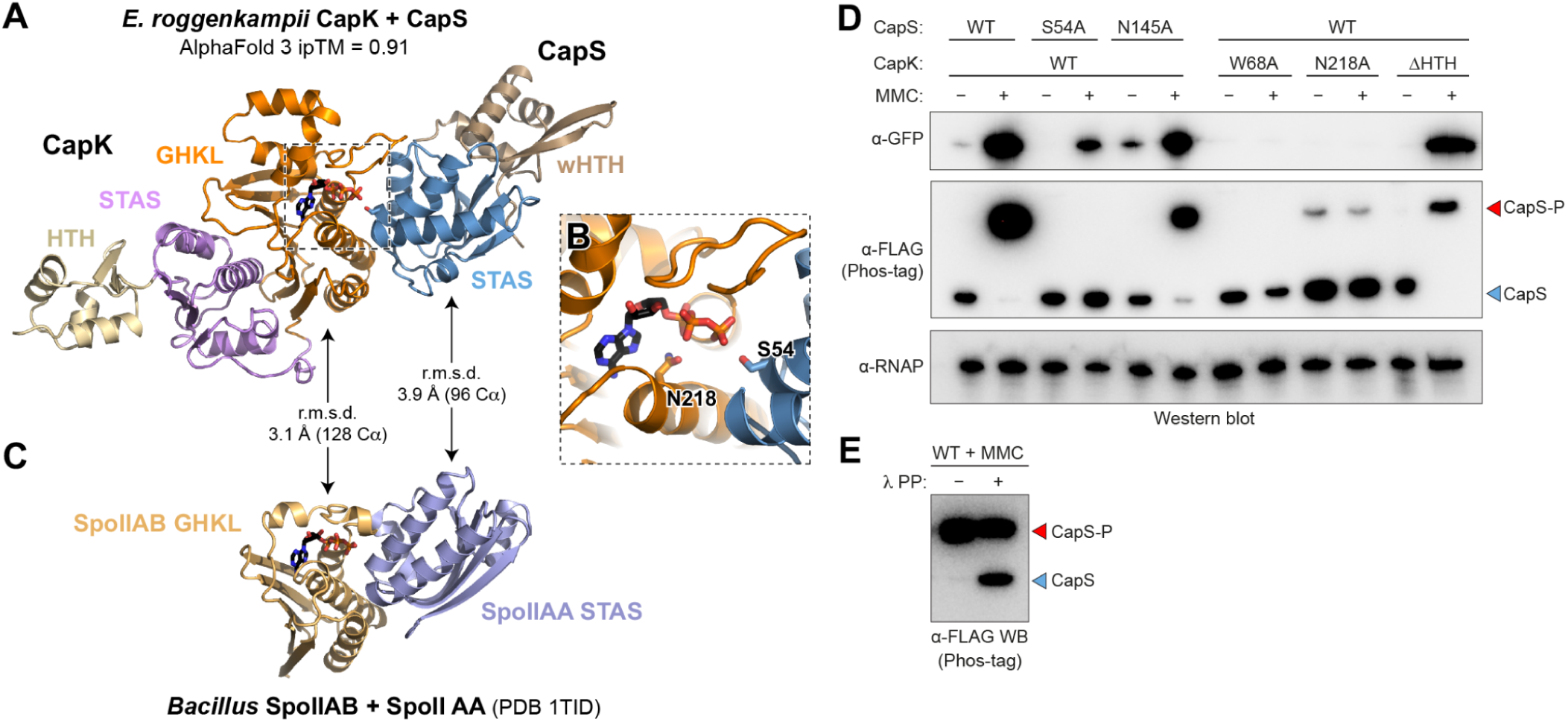
CapK phosphorylates CapS at a conserved serine residue. (A) AlphaFold 3 predicted structure of *E. roggenkampii* CapK (HTH yellow, STAS pink, GHKL orange) bound to CapS (STAS blue, wHTH brown) and ATP (black sticks). See **Figure S4B-C** for AlphaFold confidence (pLDDT and PAE). (B) Closeup of the CapK GHKL kinase active site, showing the conserved CapK active site residue Asn145 and CapS Ser54 as sticks. (C) Crystal structure of the *Bacillus* SpoIIAB-SpoIIAA complex (PDB ID 1TID) (Masuda *et al*, 2004), aligned to CapK-CapS, with Cα r.m.s.d. values of the SpoIIAB and CapK GHKL domains, and the SpoIIAA and CapS STAS domains shown. ATP bound to the SpoIIAB GHKL domain is shown in sticks. (D) Western blots showing GFP expression (α-GFP, top) and CapS mobility shift (α-FLAG Phos-tag, middle) upon addition of mitomycin C (MMC) in the GFP expression reporter expressing FLAG-CapS. Unshifted FLAG-CapS is marked “CapS” and shifted FLAG-CapS is marked “CapS-P” for phosphorylation. CapK ΔHTH is a truncated construct missing the N-terminal HTH domain. α-RNAP: anti-RpoS RNA polymerase western blot (loading control). (E) Anti-FLAG western blot of CapS isolated from cells with the wild-type GFP expression reporter exposed to mitomyin C (MMC), then treated with lambda protein phosphatase (λ PP) before loading on the Phos-tag gel.

The similarity of CapK and CapS to anti-sigma factors and their antagonists, plus our AlphaFold prediction of the CapK-CapS complex, suggested that CapK phosphorylates CapS to control transcription of the downstream operon. To test this idea, we used Phos-tag SDS-PAGE gels and detected a mobility shift of FLAG-CapS in our expression reporter upon addition of MMC, correlated with expression of GFP (**Figure 4D**). Supporting the idea that this mobility shift is due to CapS phosphorylation, treatment of samples with lambda protein phosphatase (λ PP) reversed the mobility shift (**Figure 4E**). Mutation of CapS Ser54 to alanine (S54A) prevented CapS phosphorylation, and strongly reduced MMC-dependent GFP expression (**Figure 4D**). Separately, mutation of a highly conserved asparagine residue in CapK’s kinase active site (N218A; **Figure 4B**) strongly reduced CapS phosphorylation, and prevented MMC-dependent GFP expression (**Figure 4D**). Thus, CapK phosphorylates CapS at a conserved serine upon DNA damage, dissociating the CapS homodimer and disrupting CapS-DNA binding, to mediate increased expression of the downstream operon. We note that CapS levels also increase upon MMC treatment (**Figure 4D** lanes 1-2), consistent with our ChIP-Seq data that suggests CapS regulates both its own expression and that of the downstream operon.

### CapK is activated by single-stranded DNA

Our data show that CapK+CapS de-represses transcription of the downstream operon when cells are treated with the DNA damaging agent mitomycin C (MMC), and that de-repression requires UvrA and UvrB, proteins that act early in the NER DNA repair pathway. These data suggest that CapK’s kinase activity is activated by a byproduct of DNA repair. In prior work, we found that the transcriptional regulators CapW and CapP, which also respond to DNA damage, directly bind single-stranded DNA (ssDNA) for activation (Lau *et al*, 2022; Blankenchip & Corbett, 2024). To test whether CapK is similarly activated by ssDNA, we separately purified *E. roggenkampii* CapS and CapK and incubated the two proteins with ATP and either double-stranded or single-stranded DNA. We detected a mobility shift of CapS in Phos-tag gels that depended on CapK, ATP, and single-stranded DNA (**Figure 5A**). This mobility shift was not observed when CapS Ser54 was mutated to alanine (S54A), confirming that the observed mobility shift is due to phosphorylation of Ser54 (**Figure 5A**).

**Figure 5.**
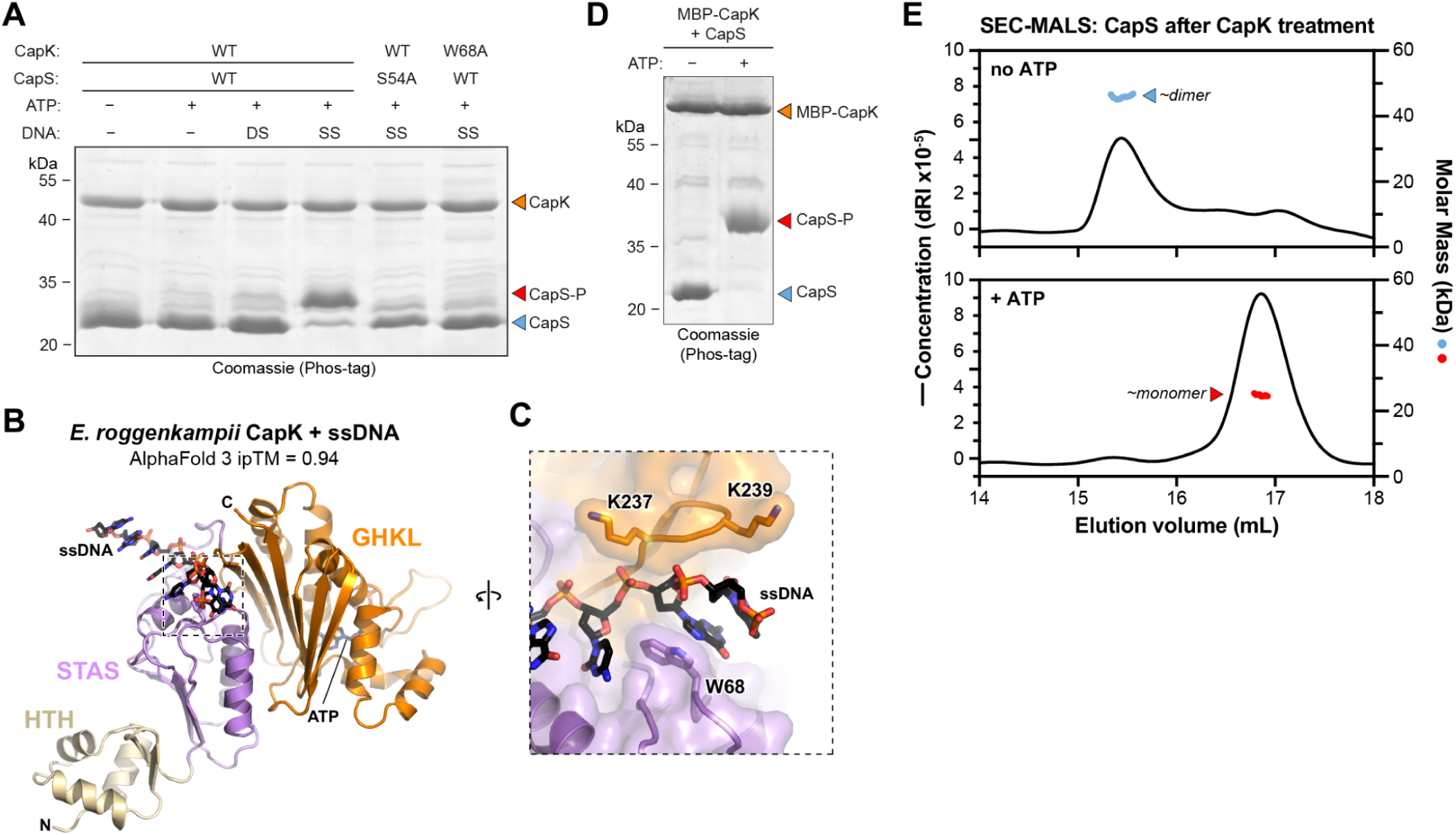
CapK is activated by ssDNA. (A) Phos-tag SDS-PAGE gel showing mobility shift of CapS upon treatment with CapK in the presence of ATP and single-stranded DNA. DS: double-stranded DNA; SS: single-stranded DNA. (B) AlphaFold 3 predicted structure of *E. roggenkampii* CapK (HTH domain yellow, STAS domain pink, GHKL domain orange) bound to ssDNA (black). See **Figure S5** for AlphaFold 3 confidence (pLDDT and PAE). (C) Closeup view of the predicted interaction between CapK and ssDNA, with CapK W68, K237, and K239 shown as sticks. (D) Phos-tag SDS-PAGE gel of CapS samples used for SEC-MALS analysis in panel (E). (E) SEC-MALS of CapS after treatment with MBP-tagged CapK and single-stranded DNA, in the absence of ATP (top) or the presence of ATP (bottom). Phosphorylated CapS shows a molecular weight consistent with a monomer (19.5 kDa), and unphosphorylated CapS shows a molecular weight consistent with a dimer (39 kDa).

We were unable to detect binding of CapK to ssDNA using a fluorescence polarization assay (not shown), so we performed an AlphaFold 3 structure prediction of *E. roggenkampii* CapK with a short poly-T ssDNA. The prediction shows a confident interaction (ipTM = 0.94) with ssDNA positioned in a positively-charged groove between the CapK STAS and GHKL kinase domains (**Figure 5B-C, S5A-D**). We identified a highly-conserved tryptophan residue (Trp68) predicted to form pi-stacking interactions with the bound DNA (**Figure 5C**). Mutation of Trp68 to alanine (W68A) eliminated ssDNA-stimulated kinase activity (**Figure 5A**), supporting the AlphaFold model for CapK-ssDNA binding. The CapK W68A mutant also disrupted both CapS phosphorylation and GFP expression in our expression reporter (**Figure 4D**).

To directly test whether phosphorylation disrupts the CapS dimer, we incubated wild-type *E. roggenkampii* CapS with MBP-tagged CapK, ATP, and ssDNA, then analyzed the reaction mixtures by both Phos-tag SDS-PAGE and SEC-MALS. We found that in the absence of ATP, CapS was not phosphorylated and was dimeric (**Figure 5D-E**). In contrast, addition of ATP and ssDNA resulted in CapS phosphorylation and complete conversion to a monomeric state (**Figure 5D-E**).

### Identification of a toxin-antitoxin locus with CapK and CapS-like proteins

In BLAST searches for CapK-related proteins in bacteria, we identified a family of proteins containing CapK-like STAS and GHKL kinase domains. Gene neighborhood analysis using webFLAGS (Saha *et al*, 2021) and *fast.genomics* (Price & Arkin, 2024) revealed that this protein is reproducibly found in a three-gene operon with an unknown protein containing a STAS domain and a short C-terminal region, and a VapC-like protein containing a predicted pilT N-terminal (PIN) nuclease domain (**Figure 6A**). VapC is the toxin component of VapBC toxin-antitoxin (TA) systems, and cleaves RNAs when it is released from a complex with its antitoxin VapB (Pandey & Gerdes, 2005; Arcus *et al*, 2011). We hypothesized that this three-gene operon possesses a similar TA-like function, and named the operon *vapKSC*, encoding a CapK-like kinase (VapK), a STAS domain protein (VapS), and a VapC-like predicted nuclease. Through iterative BLAST searches in IMG, we identified 289 instances of *vapKSC* across diverse bacterial groups, and analysis of their gene neighborhoods using PADLOC (Payne *et al*, 2022) revealed that 103 (36%) are located within 10 kb of at least one other identified anti-phage immune gene (**Table S3**).

**Figure 6.**
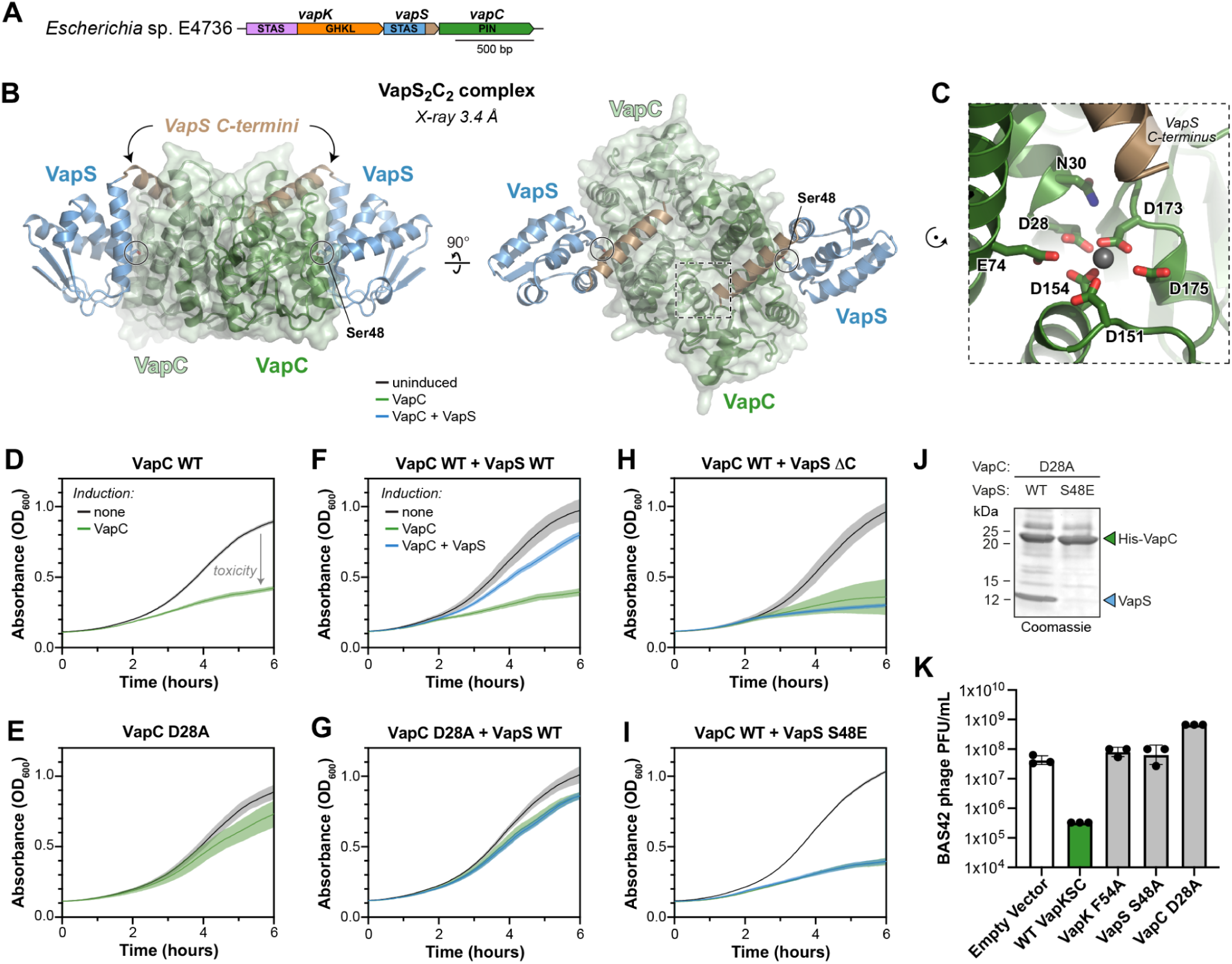
Identification of *vapKSC* operons. (A) Operon schematic of a *vapKSC* operon from *Escherichia* sp. E4736, with predicted domains of each protein noted. (B) Two views of the X-ray crystal structure of the *Escherichia* VapS_2_VapC_2_ complex, with VapS STAS domain colored blue and C-terminal region brown, and the VapC dimer colored dark green/light green. (C) Closeup of the VapC active site, with residues conserved across 279 VapC proteins in *vapKSC* operons shown as sticks, a bound Mg^2+^ ion shown in gray, and the nearby VapS C-terminus shown in brown. (D) Growth curves for *E. coli* cells transformed with an arabinose-inducible plasmid encoding *Escherichia* VapC, in non-inducing conditions (LB + 0.2% glucose, black line) or inducing conditions (LB + 0.2% arabinose, green line). Except where indicated, solid lines in panels D-I represent the average of three replicates, and shaded area indicates standard deviation. (E) Growth curves for *E. coli* cells transformed with an arabinose-inducible plasmid encoding *Escherichia* VapC D28A. The induced curve (green) represents data from duplicate samples. (F) Growth curves for *E. coli* cells transformed with an arabinose-inducible plasmid encoding *Escherichia* VapC plus an IPTG-inducible plasmid encoding VapS, in non-inducing conditions (LB + 0.2% glucose, black line), VapC-inducing conditions (LB + 0.2% arabinose, green line), and VapC+VapS-inducing conditions (LB + 0.2% arabinose + 50 µM IPTG, blue line). The uninduced curve (black) represents data from duplicate samples. (G) Growth curves for *E. coli* cells transformed with an arabinose-inducible plasmid encoding *Escherichia* VapC D28A plus an IPTG-inducible plasmid encoding VapS. (H) Growth curves for *E. coli* cells transformed with an arabinose-inducible plasmid encoding *Escherichia* VapC plus an IPTG-inducible plasmid encoding VapS ΔC. (I) Growth curves for *E. coli* cells transformed with an arabinose-inducible plasmid encoding *Escherichia* VapC plus an IPTG-inducible plasmid encoding VapS S48E. (J) SDS-PAGE gel showing results of a Ni^2+^ pulldown assay after coexpression of His_6_-tagged VapC D28A with VapS wild type (WT; left lane) or S48E (right lane). (K) Plaque assay showing VapKSC-mediated protection of *E. coli* against phage AndreasVesalius (Bas42).

Based on the function of VapBC systems and our findings with CapK and CapS, we hypothesized that VapS binds VapC and functions as an antitoxin, and that VapK phosphorylates VapS to disrupt the VapSC complex and activate VapC. Supporting this model, we were unable to express and purify a VapC protein from *Escherichia* sp. E4736 in *E. coli* due to toxicity, but we could co-express the protein with its cognate VapS. By SEC-MALS, we found that *Escherichia* VapS and VapC form a complex with 2:2 stoichiometry (**Figure S6**). We crystallized and determined a 3.4 Å resolution X-ray crystal structure of the *Escherichia* VapSC complex (**Table S5, Figure S7**), revealing a 2:2 heterotetramer with a central VapC dimer, and each VapC protomer bound to one protomer of VapS (**Figure 6B**). VapS binds VapC primarily via its STAS domain, and the VapS C-terminus forms an α-helix that extends to within ∼6 Å of the VapC catalytic site (**Figure 6B-C**). Unlike VapB proteins, many of which use a conserved arginine residue to displace the catalytic Mg^2+^ ion from the VapC active site (Miallau *et al*, 2009; Dienemann *et al*, 2011; Min *et al*, 2012; Kang *et al*, 2020), VapS does not displace Mg^2+^ from the VapC active site (**Figure 6C**). Instead, VapS may block binding of the RNA target of VapC, preventing catalysis.

Sequence alignments of VapS reveal that, like CapS, this protein possesses an absolutely conserved serine or threonine residue (*Escherichia* VapS Ser48) in its STAS domain. In our structure of the VapSC complex, this residue is buried in the VapS-VapC interface (**Figure 6B**). In an AlphaFold 3 predicted structure of a VapK-VapS complex, the conserved serine residue is positioned for phosphorylation by the VapK kinase domain, as in prior structures of bacterial anti-sigma factor antagonists and their cognate kinase anti-sigma factors (**Figure S8A-D**). To test whether VapS regulates VapC toxicity, we used a bacterial growth assay with VapC and VapS controlled by separate inducible promoters. As expected, expression of wild-type VapC but not a catalytic-dead VapC mutant (Asp28 to alanine; D28A) resulted in strong toxicity (**Figure 6D-E**). Coexpression of VapS alongside VapC suppressed VapC-mediated toxicity, confirming that VapS is an antitoxin (**Figure 6F-G**). Deletion of the VapS C-terminal region (residues 94-116) rendered the protein unable to counteract VapC toxicity, confirming that VapS directly inhibits VapC activity through its C-terminus (**Figure 6H**). Mimicking VapS phosphorylation with a Ser48 to glutamate (S48E) also VapS unable to counteract VapC toxicity (**Figure 6I**). Supporting this idea, when we purified VapC (catalytic mutant D28A) from *E. coli* cells also expressing either wild-type VapS or the VapS S48E mutant, we found that VapC co-purified with wild-type VapS but not the VapS S48E mutant (**Figure 6J**).

To determine the activation mechanism of VapKSC, we performed an AlphaFold 3 prediction of a VapK-ssDNA complex, resulting in a high-confidence prediction with an ipTM of 0.94, showing ssDNA bound at the interface of the STAS and GHKL kinase domains (**Figure S9A-D**). Close inspection of the predicted structure identified a conserved phenylalanine residue in VapK (Phe54) equivalent to CapK Trp68 (**Figure S9B**). These results suggest that, like CapK, VapK is activated by binding ssDNA resulting from DNA damage. We were unfortunately unable to test this idea: VapK is insoluble when expressed in *E. coli*; and we could not detect a FLAG-tagged VapS on western blots, presumably because of low expression. We did, however, find that the *vapKSC* operon provided a ∼2-log protection against at least one bacteriophage, *E. coli* phage AndreasVesalius (Bas42) (Maffei *et al*, 2021), in the *Tequatovirus* family (**Figure 6K**). Mutation of VapC (D28A), VapS (S48A), or the VapK putative ssDNA binding site (F54A) all eliminated protection against this phage (**Figure 6K**). Overall, these data support a model in which DNA damage (arising from phage infection or other stress) activates VapK, which phosphorylates VapS, releasing it from VapC.

## DISCUSSION

Here we identify CapK and CapS, which control the expression of downstream immune operons – Bub, CBASS, and others – in response to DNA damage. Our data show that CapS is a homodimeric DNA-binding transcriptional repressor that binds its own promoter and that of the downstream immune operon to suppress transcription (**Figure 7A**). Upon binding ssDNA byproducts of DNA repair, CapK is activated and phosphorylates a conserved serine in the STAS domain of CapS, disrupting the CapS dimer to de-repress immune operon transcription (**Figure 7A**). CapK+CapS is the newest example of immune operon-associated transcriptional regulators that sense and respond to DNA damage, a group that also includes CapW/BrxR (Blankenchip *et al*, 2022; Blankenchip & Corbett, 2024; Picton *et al*, 2022; Luyten *et al*, 2022) and CapH+CapP (Lau *et al*, 2022). These regulators may prevent toxicity arising from constitutive expression of immune operons, especially abortive infection systems like CBASS whose high-level activation can kill the host cell (Ye *et al*, 2020; Cohen *et al*, 2019; Lopatina *et al*, 2020). Since phage infection often results in DNA damage and the production of excess ssDNA, sensing ssDNA allows these regulators to drive increased expression of immune pathways when they are needed most. Additionally, CapK+CapS and related regulators may enable a bacterium to coordinate immune responses between front-line restriction modification pathways (which themselves generate DNA damage that could activate CapK) and later-acting abortive infection pathways (Oshiro *et al*, 2025). Supporting this idea, nearly 20% of identified *capK*+*capS* genes are encoded in a larger locus where *capK*+*capS* is sandwiched between an upstream restriction-modification operon and a downstream immune operon like CBASS or DUF4747/CarolAnn (**Figure 1A**).

**Figure 7.**
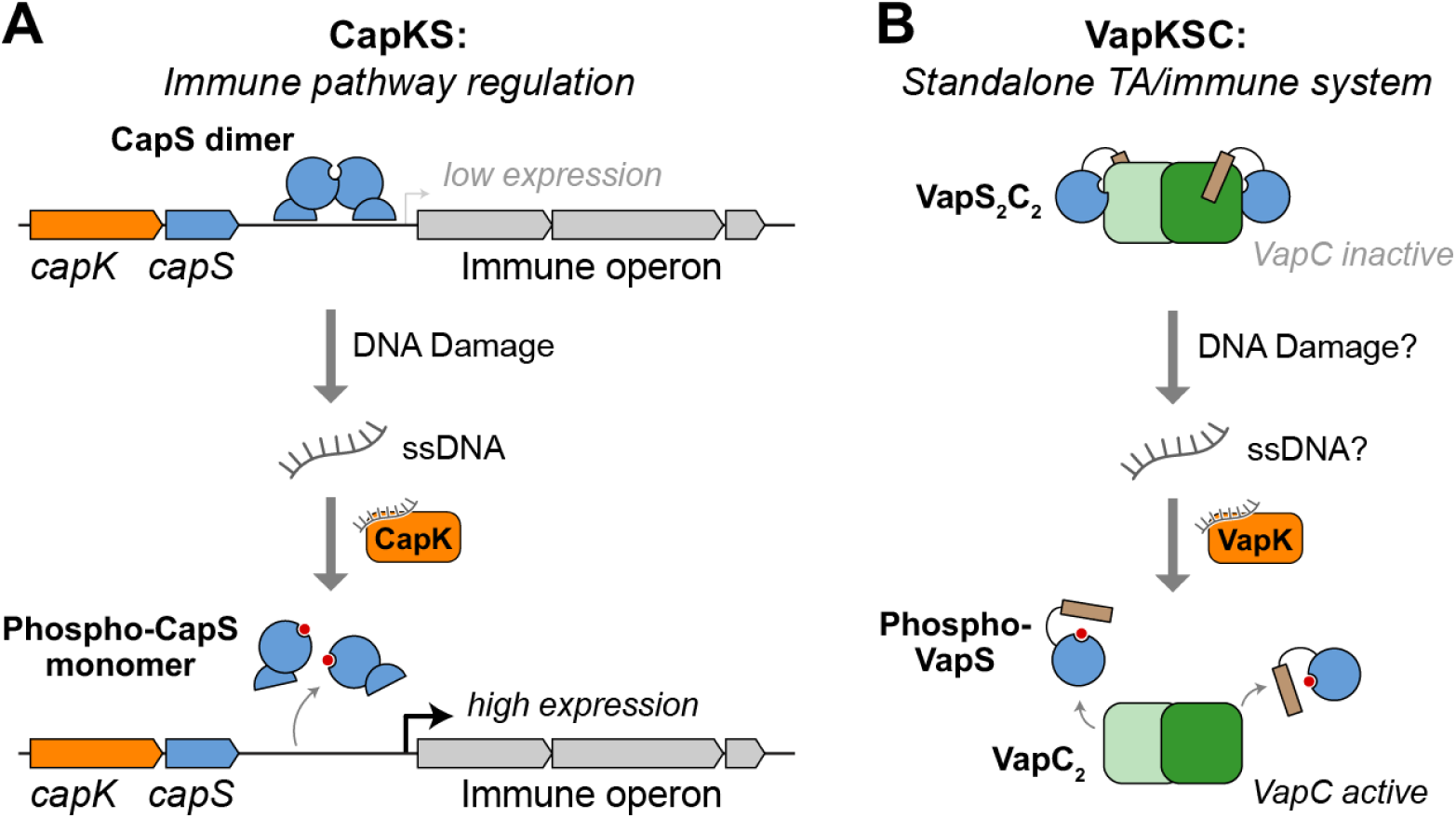
Proposed mechanism for CapKS and VapKSC. (A) Model for immune operon regulation by CapKS. In unperturbed cells, CapS forms a DNA-binding homodimer (blue) that suppresses transcription of its downstream operon (gray). Upon DNA damage, ssDNA is produced that binds and activates CapK (orange). CapK phosphorylates CapS (red circles), disrupting the CapS dimer and releasing it from DNA to allow increased expression of the downstream operon. (B) Model for VapKSC. In unperturbed cells, VapC (green) and VapS (blue/brown) form an inhibited VapC_2_S_2_ complex. Upon DNA damage and/or phage infection, ssDNA is produced that binds and activates VapK (orange). VapK phosphorylates VapS (red circles), releasing it from VapC to release inhibition of VapC.

In addition to regulating expression of the downstream immune operon, our ChIP-Seq results suggest that CapS regulates its own expression, binding its own promoter in unperturbed cells and dissociating when it is phosphorylated by CapK. Supporting the idea that CapS regulates its own expression, our reporter plasmids show increased expression of both GFP and CapS upon exposure to DNA damage. This autoregulatory circuit likely maintains CapS at levels that are (1) high enough to suppress both its own transcription and that of the downstream operon, but (2) low enough that activated CapK can readily phosphorylate the existing pool of CapS to release it from DNA. Strict control over CapS protein levels would therefore maintain the system in a poised state that is maximally sensitive to DNA damage arising from phage infection and/or activation of DNA-targeting immune pathways.

We find that CapS can bind both “wide” and “narrow” palindromic sequences with equivalent half-sites but significantly different spacing. The CapS homodimer that exists in solution is assembled through dimer interface 1, with the two CapS protomers’ wHTH domains perfectly positioned to bind a “wide” palindromic site. In contrast, “narrow” palindromic sites match the spacing between wHTH domains of CapS protomers assembled through interface 2. Our ChIP-Seq data suggests that CapS phosphorylation, which disrupts dimer interface 1, unexpectedly disrupts CapS binding to both wide and narrow palindromes. Together, this data suggests that narrow palindrome binding involves two CapS interface-1 dimers binding neighboring sites along DNA and associating with one another through dimer interface 2. This difference in recognition mode between CapS binding sites in the downstream operon’s promoter (wide palindrome) and the *capS* promoter (narrow palindrome) means that the binding affinity for these sites could be independently tuned through mutations in the two CapS dimer interfaces, thereby enabling CapS to differentially regulate its own expression and that of the downstream operon. Testing this hypothesis will require further study, including isolation of a CapS mutant that specifically disrupts dimer interface 2.

Alongside CapK+CapS, we identify a related toxin/antitoxin system we term VapKSC, in which a CapS-like protein (VapS) binds and inhibits a VapC-family toxin (**Figure 7B**). Our data suggest that the CapK-like kinase VapK is activated by ssDNA arising from DNA damage and/or phage infection. VapK-mediated phosphorylation of VapS causes dissociation of the VapC-VapS complex, resulting in activation of VapC and inhibition of cell growth, likely through cleavage of ribosomal RNA (rRNA) or tRNA. Our finding that a GHKL kinase + STAS domain protein pair function in multiple contexts to regulate different biochemical activities in response to a common signal reflects the emerging theme of modularity in the evolution and function of bacterial immune pathways. As the number of identified immune pathways has expanded into the hundreds, diverse immune pathways have been shown to share common sensor domains, effector domains, and activation mechanisms (Mariano & Blower, 2023; Georjon & Bernheim, 2023). This modularity likely enables bacteria to swiftly respond to phages that evolve to evade specific immune pathways’ detection or effector mechanisms.

We identify *capK*+*capS* adjacent to several known anti-phage immune pathways including CBASS, Thoeris, Bil, and restriction-modification systems (**Figure 1B**). The majority of *capK*+*capS* genes we identify are encoded adjacent to Bub operons, which can also be regulated by CapW or CapH+CapP, but have not been demonstrated to protect a host cell against phage infection (Ye *et al*, 2025). Bub operons, as well as other *capK*+*capS* associated genes including *yegP*, may be anti-phage immune pathways or may instead mediate more global stress responses. Indeed, the line between anti-phage immune pathways and stress-response pathways is likely fuzzier than generally appreciated, with many pathways - like VapKSC and potentially other toxin-antitoxin systems - protecting their host against phage infection as a result of their response to a global stress signal like DNA damage.

## MATERIALS AND METHODS

### Bioinformatics

Initial identification and annotation of *capK* and *capS* genes associated with Bub operons (Ye *et al*, 2025) was performed by manual inspection of gene neighborhoods using the Integrated Microbial Genomes & Microbiomes server (IMG: https://img.jgi.doe.gov) (**Table S1**). For comprehensive identification of *capK*+*capS* associated immune operons, representative CapK proteins were used as queries for exhaustive BLAST searches in IMG, followed by manual annotation of CapS-encoding genes (**Table S2**). Genomic DNA sequences +/- 10 kb of each *capKS* locus was downloaded from IMG and submitted to PADLOC (Payne *et al*, 2022) for identification of nearby immune operons. Gene neighborhoods were further annotated by manual inspection in IMG.

For identification of *vapKSC* operons, representative CapK homologs lacking the N-terminal HTH domain were used for exhaustive BLAST searches in IMG. Genomic DNA sequences +/-10 kb of each *vapK* gene was downloaded from IMG and searched for *vapC* genes using tblastn in BLAST+ (installed locally). The same sequences were submitted to PADLOC for identification of nearby immune operons. Hits were manually annotated in IMG to identify all cases containing *vapK*, *vapS*, and *vapC* (**Table S3**).

### Cloning, expression, and protein purification

Sequences of all proteins used in this study are in **Table S4**. To generate the GFP reporter plasmid, the native sequence of a CapK+CapS operon and the associated system’s promoter from *Enterobacter roggenkampii* strain EB00176-1 (reverse complement of bases 880,520-882,344 of NCBI RefSeq NZ_CM130038.1) followed by a gene encoding GFP was synthesized (IDT) and cloned via isothermal assembly into pTRC99a, which encodes an IPTG-inducible promoter. Subsequently, an N-terminal FLAG epitope tag was introduced on *capS* using PCR mutagenesis. For ChIP-Seq, a longer insert including the entire intergenic region upstream of *capK* was cloned into pTRC99a (reverse complement of bases 880,520-883,004 of NCBI RefSeq NZ_CM130038.1) followed by a gene encoding GFP.

For protein expression, *E. roggenkampii* CapK (NCBI WP_001567866.1) and CapS (NCBI WP_001567865.1) were amplified by PCR and cloned into UC Berkeley Macrolab vectors 2B-T (Addgene #29666; encoding an N-terminal TEV protease-cleavable His_6_-tag), or 13S-A (Addgene #48323) for expression and purification. Point mutations were generated by PCR-based mutagenesis. Genes encoding full length CapK (NCBI WP_160230705.1) and CapS (NCBI WP_046127252.1) from *V. cholerae* strain 964015 (genome sequence NCBI RefSeq NZ_JAWJBE020000004.1) were codon-optimized for *E. coli* expression, synthesized (IDT) and cloned into UC Berkeley Macrolab vector 2B-T or 13S-A.

To generate the VapKSC plasmid used for bacteriophage infection assays, the native sequence of a VapK+VapS+VapC operon from *Escherichia* sp. E4736 (bases 78125-80342 of NCBI RefSeq NZ_VATT01000011.1) was cloned into vector pTRC99a (Amann *et al*, 1988). *Escherichia* sp. E4736 VapK (NCBI WP_135559716.1), VapS (NCBI WP_135559717.1), and VapC (NCBI WP_135559718.1) were cloned into UC Berkeley Macrolab vectors 2B-T or 13S-A for expression and purification. For toxicity assays, VapC was cloned into pBAD LIC cloning vector (8A) (Addgene #37501; no tag), and VapS was cloned into UC Berkeley Macrolab vector 13S-A. Point mutations were generated by PCR-based mutagenesis.

### GFP expression reporter assays

For GFP reporter expression assays, *E. coli* MG1655 cells were transformed with GFP expression reporter plasmids (wild type or indicated mutants), and 2 mL cultures in LB media plus 100 µg/mL ampicillin were grown overnight at 37°C with shaking. The next morning, 50 µL of saturated overnight culture was diluted into 5 mL fresh media and grown to an OD_600_ of 0.3. Cells were stressed by addition of 10 µg/mL mitomycin C (MMC), 10 µg/mL levofloxacin, 10 µg/mL colistin, or 100 µg/mL zeocin and grown for a further 1 hour at 37°C. For GFP expression analysis, 500 µL was centrifuged, then the cell pellet was resuspended in 50 µL 2X SDS-PAGE loading buffer (125 mM Tris-HCl pH 6.8, 20% Glycerol, 4% SDS, 200 mM DTT, 0.01% w/v bromophenol blue). Sample volumes were adjusted based on the culture density. Samples were placed in a boiling water bath for 3 minutes, then 5 µL samples were loaded onto a 4–20% Mini-PROTEAN TGX gel (Bio-Rad, 4561096) or 10% SDS-PAGE gel supplemented with 50 µM Phos binding reagent (APExBIO F4002) and separated at 180 V for 40 minutes at room temperature or 4°C, respectively.

For GFP expression reporter assays in single-gene knockout strains, single-gene knockout *E. coli* strains from the KEIO collection (Baba *et al*, 2006) were obtained from Horizon Discovery (Strain IDs: parent strain BW25113 (Datsenko & Wanner, 2000); *ΔrecA* JW2669, *ΔrecB* JW2788, *ΔuvrA* JW4019, *ΔuvrB* JW0762, *ΔuvrC* JW1898, *ΔuvrD* JW3786).

### Western blots

For western blots, samples were separated on 4–20% Mini-PROTEAN TGX gels (Bio-Rad, 4561096) and then transferred to PVDF membranes (Bio-Rad Turbo transfer kit, 170-4272). PVDF membranes (Bio-Rad) were blocked for 1 hour at room temperature with 5% nonfat dry milk in TBST (50 mM Tris-HCl pH 8.5, 150 mM NaCl, 0.1% Tween-20) followed by blotting for primary antibody overnight at 4°C or room temperature for 1 hour (Antibodies used: Mouse anti-GFP primary antibody (Roche, 11814460001) at 1:3000 dilution; Mouse anti-FLAG primary antibody (Sigma-Aldrich, F3165) at 1:5000 dilution; Mouse anti-RNA polymerase primary antibody (clone NT63; BioLegend #10019-878) at 1:3000 dilution). Blots were washed three times in TBST and incubated with Goat anti-mouse HRP-linked secondary antibody (Millipore Sigma, 12-349) at dilution 1:4000. Membranes were washed three times in TBST, incubated with ECL detection reagent (Cytiva RPN2232) for 1 minute, and imaged on a Bio-Rad ChemiDoc imager.

### Protein Expression and Purification

Proteins were expressed in *E. coli* strain Rosetta 2 (DE3) pLysS (EMD millipore). Cultures were grown in 1L of 2XYT medium at 37°C to A_600_ = 0.6, induced with 0.25 mM IPTG and moved to 20°C for 16-18 h. Cells were harvested by centrifugation and resuspended in buffer A (25 mM Tris pH 8.5, 10% glycerol, 300 mM NaCl, 5 mM MgCl_2_ and 1 mM NaN_3_) plus 5 mM imidazole and 5 mM β-mercaptoethanol and lysed by sonication (Branson sonifier). Lysates were clarified by centrifugation, then passed over Ni-NTA agarose (Qiagen) in resuspension buffer, washed in wash buffer (buffer A containing 20 mM imidazole), and eluted in elution buffer (buffer A containing 400 mM imidazole). Proteins were then concentrated by ultrafiltration (Amicon Ultra; EMD millipore). All proteins were passed over a Superdex 200 Increase size exclusion column (Cytiva) in buffer A containing 1 mM DTT. Peak fractions were concentrated by ultrafiltration and stored at 4°C.

### Crystallization and structure determination

For crystallization of *E. roggenkampii* CapS, purified protein at 9 mg/mL in crystallization buffer (25 mM Tris pH 8.5, 0.2 M NaCl, 5 mM MgCl_2_, and 1 mM TCEP) was mixed 1:1 with well solution containing 0.2 M Ammonium sulfate and 13% (w/v) PEG 4000 in hanging drop format. Crystals were collected into a cryoprotectant solution containing an additional 15% glycerol and frozen in liquid nitrogen. Diffraction data were collected at beamline 12-1 at the Stanford Synchrotron Light Source (SSRL), and processed with the autoPROC pipeline which uses XDS (Kabsch, 2010) for indexing and integration, AIMLESS (Evans & Murshudov, 2013)for merging, and CTruncate (Evans & Murshudov, 2013) for conversion to structure factors. The structure was determined by molecular replacement in PHASER (McCoy *et al*, 2007) using an AlphaFold 3-predicted structure, manually rebuilt in COOT (Emsley *et al*, 2010), and refined in phenix.refine (Adams *et al*, 2010) using individual atomic position and isotropic B-factor refinement, plus TLS refinement (one TLS group per chain).

For crystallization of *V. cholerae* CapS (form 1), copurified CapK and CapS (S58A mutant) at 14 mg/mL plus 2 mM ATP in crystallization buffer (25 mM Tris pH 8.5, 0.2 M NaCl, 5 mM MgCl_2_, and 1 mM TCEP) was mixed 1:1 with well solution containing 0.2 M MgCl_2_, 0.1 M Bis-Tris pH 5.5, and 20% (w/v) PEG 3350 in hanging drop format. Crystals were collected into a cryoprotectant solution containing an additional 10% glycerol and frozen in liquid nitrogen. For crystallization of *V. cholerae* CapS (form 2), copurified CapK and CapS (S58A mutant) at 14 mg/mL plus 2 mM ATP in crystallization buffer (25 mM Tris pH 8.5, 0.2 M NaCl, 5 mM MgCl_2_, and 1 mM TCEP) was mixed 1:1 with well solution containing 0.2 M Ammonium sulfate, 0.1 M sodium acetate pH 4.6, and 17% (w/v) PEG 4000 in hanging drop format. Crystals were collected into a cryoprotectant solution containing an additional 10% glycerol and frozen in liquid nitrogen. For both form 1 and form 2, diffraction data were collected at SSRL beamline 9-2, processed with the autoPROC pipeline, and the structures were determined by molecular replacement in PHASER using AlphaFold 3-predicted structures. Models were manually rebuilt in COOT and refined in phenix.refine using individual atomic position and isotropic B-factor refinement, plus TLS refinement (one TLS group per chain).

For crystallization of the *Escherichia* VapS-VapC complex, copurified VapS+VapC at 15 mg/mL in crystallization buffer (25 mM Tris pH 8.5, 0.2 M NaCl, 5 mM MgCl_2_, and 1 mM TCEP) was mixed 1:1 with well solution containing 0.1 M HEPES pH 7.5, 10% PEG 8000, and 0.2 M Calcium Acetate in sitting drop format. Crystals were collected into cryoprotectant solution containing an additional 20% glycerol and frozen in liquid nitrogen. Diffraction data were collected at the Advanced Photon Source at Argonne National Laboratory, on beamline 24ID-C. Data were indexed and integrated with XDS (Kabsch, 2010) in the RAPD2 pipeline (https://git.nec.aps.anl.gov/rapd/rapd). Because of high anisotropy, unmerged data were uploaded to the STARANISO web server (https://staraniso.globalphasing.org/) for application of an ellipsoidal resolution cutoffs (4.46 Å along a* and b*; 3.4 Å along c*) and anisotopy-corrected scaling. Values in **Table S5** are after STARANISO processing and merging. Graphs showing merging statistics by resolution (completeness, multiplicity, I/σ, *R_merge_*/*R_sym_*/*R_pim_*, and CC_1/2_) are shown in **Figure S7**. The structure was determined by molecular replacement in PHASER (McCoy *et al*, 2007) using an AlphaFold 3 model of a 2:2 VapS-VapC complex (2 heterotetramers placed). The N-terminal tag for two VapS protomers, plus Mg^2+^ ions in each VapC active site, were manually built in COOT (Emsley *et al*, 2010). The structure was refined to a resolution of 3.4 Å in phenix.refine (Adams *et al*, 2010) with reference model restraints (using the input AlphaFold 3 model of VapS-VapC), strict non-crystallographic symmetry constraints, and grouped B-factors (two groups per residue).

Raw diffraction data and refined structure factors & models have been deposited at the SBGrid Data Bank (https://data.sbgrid.org) and the RSCB Protein Databank (https://rcsb.org), respectively (accession numbers in **Table S5**).

### SEC-MALS

For characterization of CapS oligomeric states by size exclusion chromatography coupled to multi-angle light scattering (SEC-MALS), 100 µL of purified protein at a concentration of 2 mg/mL was injected onto a size exclusion column (Superdex 200 Increase 10/300 GL, Cytiva) in SEC-MALS buffer (25 mM Tris pH 8.5, 300 mM NaCl, 5 mM MgCl_2_, 1 mM NaN_3_, 1 mM DTT, and 5% glycerol). Light scattering and differential refractive index (dRI) profiles were collected using miniDAWN TREOS and Optilab T-rEX detectors (Wyatt Technology). SEC-MALS data were analyzed using ASTRA software version 8 and visualized with Graphpad Prism v.10.1.1.

### DNA binding assays

For characterization of DNA binding by fluorescence polarization assays, double stranded DNA oligos were produced by annealing 24 bp complementary oligos (sequences highlighted in **Figure 2C**), one of which was 5’ 6-FAM labeled. FP reactions (30 µL) in buffer 25 mM Tris pH 8.5, 50 mM sodium glutamate, 5 mM MgCl_2_, 1 mM DTT, 5% glycerol, and 0.01% NP40 analog contained 50 nM DNA and the indicated protein concentration. FP was read using a Tecan Spark plate reader, and binding data were analyzed with Graphpad Prism v.10.1.1 using a single-site binding model.

### CapK kinase assays

For detecting phosphorylation of CapS by CapK in cells, DNA damage assays were performed with 10 µg/mL mitomycin C and the *E. roggenkampii* GFP reporter system with a FLAG-tag fused to the N-terminus of CapS. The resuspended cell pellet in 2x SDS-PAGE loading buffer was boiled for 3 minutes, and 5 µL of sample was loaded onto a 10% SDS-PAGE gel containing 50 uM of Phos binding reagent (APExBIO F4002). Gels were run at 180V for 40 minutes at 4°C and washed twice with 5 mM EDTA for 10 min before western blotting.

For detection of CapK kinase activity *in vitro*, 20 µL reactions containing 10 µM purified *E. roggenkampii* CapK and CapS, 2 mM ATP, and 10 uM ssDNA in reaction buffer (20 mM HEPES pH 7.5, 100 mM NaCl, 20 mM MgCl_2_, 1 mM DTT) were incubated 1 hour at 37°C, then added to 20 µL 2xSDS sample buffer. 10 µL of each sample was loaded onto a 10% SDS-PAGE gel containing 50 uM of Phos binding reagent (APExBIO F4002) and Coomassie-stained for visualization.

### Chromatin immunoprecipitation (ChIP) and sequencing

For chromatin immunoprecipitation experiments, *E. coli* MG1655 cells transformed with the *capK-capS* region plasmid (wild-type or *capS*-S54E) were grown from fresh streaks in LB medium with ampicillin, inoculating 10 mL secondary cultures with 100 μL overnight culture and growing to an OD₆₀₀ of ∼0.3. Where indicated, DNA damage was induced by addition of mitomycin C to a final concentration of 5 μg/mL followed by incubation for 1 hour; untreated controls were processed in parallel. Cultures were crosslinked by addition of molecular-grade formaldehyde to a final concentration of ∼1% and incubated for 10 min at 37°C with shaking, followed by quenching with glycine (final concentration 125 mM) for 10 min. Cells were pelleted, washed twice with ice-cold PBS, and stored at −80°C until processing.

Frozen pellets were thawed on ice and lysed in cell lysis buffer (50 mM HEPES pH 8.0, 1 mM EDTA, 85 mM KCl, 0.5% NP40) supplemented with protease inhibitors. Chromatin was transferred to Covaris microTUBEs and sheared to an average fragment size of ∼200 bp. After sonication, samples were clarified by centrifugation to remove debris, and an aliquot of input chromatin was de-crosslinked, proteinase K-treated, and purified to assess fragment size using an Agilent TapeStation (D1000). For immunoprecipitation, chromatin was diluted in dilution buffer (0.1% SDS, 1.1% Triton X-100, 1.2 mM EDTA, 165 mM NaCl, 16.7 mM Tris-HCl pH 8.1) and incubated with antibody-conjugated magnetic beads for 2 h at 4°C with rotation; mock IPs were performed using non-specific beads. Beads were sequentially washed with low-salt buffer (Tris•HCl 50 mM, pH 8.0, 150 mM NaCl, 0.1% SDS, 1% NP40, 1 mM EDTA and 0.5% Deoxycholate Na), high-salt buffer (Tris•HCl 50 mM, pH 8.0, 500 mM NaCl, 0.1% SDS, 1% NP40, 1 mM EDTA, 0.5% Deoxycholate Na), LiCl buffer (Tris•HCl 50 mM, pH 8.0 250 mM LiCl 0.1% SDS 1% NP40 1 mM EDTA 0.5% Deoxycholate Na), and TE buffer (Tris•HCl 10 mM, pH 8.0, 0.25 mM EDTA), followed by elution in 1% SDS, 100 mM NaHCO₃ at 65°C. Crosslinks were reversed by overnight incubation at 65°C, samples were treated with proteinase K, and DNA was purified using Zymo columns (D4004).

ChIP and Mock DNA were converted into sequencing libraries using NEBNext Ultra II DNA Library Prep reagents with indexed adapters, followed by PCR amplification. Library quality and fragment size distributions were assessed on an Agilent TapeStation 4150. Libraries were pooled and sequenced on an Illumina NovaSeq platform using paired-end 150 bp reads, yielding >20 million reads per sample. Sequencing quality was evaluated using FastQC prior to downstream analysis.

Three independent biological replicates for each sample were sequenced. High quality reads were selected for further analysis. The reads were aligned to the plasmid sequence using bowtie 2/2.5.0 (Langmead *et al*, 2009; Langmead & Salzberg, 2012). PCR duplicates were removed following best practices for paired-end data using samtools fixmate and markdup (Danecek *et al*, 2021; Li *et al*, 2009). Only primary, properly mapped read pairs were retained for downstream analyses. Genome-wide coverage tracks were generated from deduplicated BAM files using bedtools genomecov, normalized to reads per million mapped reads (RPM), and subsequently aggregated into fixed-width bins (10–20 bp) to reduce local noise on the small plasmid genome. Because ChIP-seq signal is strand-independent, all coverage tracks were generated without strand filtering. Enrichment was calculated as the log2 ratio of IP over Mock RPM-normalized coverage, with the addition of a small pseudocount to avoid division by zero. For each replicate, putative binding regions were identified using a percentile-based approach, defining enriched bins as those within the top 0.5-1% of genome-wide log2 enrichment values, rather than applying a fixed fold-enrichment cutoff. Adjacent enriched bins were merged if separated by ≤40 bp, and only regions ≥40 bp in length were retained. To ensure robustness, enrichment profiles were assessed independently for each biological replicate, and only regions reproducibly detected in at least two replicates per condition were considered high-confidence binding sites. All coverage and enrichment tracks, as well as reproducible peak intervals, were visualized in the Integrative Genomics Viewer (IGV) using consistent genomic coordinates and fixed y-axis scales to enable direct comparison across conditions.

Sequencing data have been deposited at the NCBI Gene Expression Omnibus (GEO) database under BioProject ID PRJNA1415371.

### Toxicity assays

For bacterial growth curves, overnight cultures of *E. coli* strain SoluBL21 (Genlantis) with plasmids of interest were grown at 37°C in LB plus appropriate antibiotics. Cultures were diluted OD_600_=0.1 in fresh LB with antibiotics and grown at 37°C until OD_600_=0.5. The cells were diluted back to OD_600_=0.1 with LB plus antibiotics and the inducers (0.2% arabinose, 50 μM IPTG). 100 μL of diluted cultures were plated to standard clear 96 well plate with lid (Corning) and incubated in a plate reader (Tecan Spark) at 37°C. The OD600 of each well was measured every 3 minutes for 8 hours total with medium intensity shaking. Each sample included three independent replicates on a single plate.

For toxicity spotting assays, overnight cultures of *E. coli* strain MG1655 with plasmids of interest were grown at 37°C in LB plus appropriate antibiotics. Cultures were diluted OD_600_=0.1 in fresh LB with antibiotics supplemented with 0.2% glucose to prevent toxicity and grown at 37°C until OD_600_=0.5. Tenfold serial dilutions of these cultures were prepared in LB broth, and 3 μl of these dilutions was spotted onto LB agar plates containing the appropriate antibiotic and inducer (0.2% arabinose) or suppressor (0.2% glucose). The plates were incubated overnight at 37°C and were imaged the next day using a ChemiDoc Imaging System (Bio-Rad).

### Bacteriophage infectivity

Plaque-Forming Unit (PFU) estimations were used as a measure of phage infections. Briefly, M9 minimal media agar plates were prepared with 10 μM IPTG and left to dry overnight at room temperature. Five overnight starter cultures of *E. coli* ΔRM (Alexander Harms) (Maffei *et al*, 2021) harboring either the pTRC99a empty vector or *vapKSC* in pTRC99a (wild type and indicated mutants) were diluted in M9 minimal media+ampicillin and grown at 37°C until the OD_600_ reached 0.6-0.8. Next, 0.1 mL of this culture was mixed with 5 mL of preheated 0.35% top agar (composed of M9 minimal media plus 0.7% agar at a 1:1 ratio) at 55°C, and the mixture was overlaid onto M9+ampicillin agar plates. 10-fold dilutions of Bas42 phage stock was prepared in the BASEL phage buffer (0.05 M Tris pH 7.5, 0.1 M NaCl, and 10 mM MgSO_4_) as the diluent. After 1 hour of air drying of plates, 3 μL volume of 10-fold phage dilutions were spotted on these plates and allowed to absorb into the top agar for 30 minutes, followed by overnight incubation at 37°C. The plaques were quantified the next day by counting individual plaques in the highest dilution that individual plaques were visible, and the reported values are the mean plus standard deviation of three biological replicates. Images were taken using an Epson flatbed scanner.

## ACKNOWLEDGEMENTS

The authors thank members of the Corbett Lab and A. Whiteley for helpful discussions, and the UCSD Physics Computing Group for computational support. The authors acknowledge funding from the HHMI Emerging Pathogens Initiative (to E.V., and K.D.C) and the National Institutes of Health (R35 GM144121 to K.D.C, T32 GM139795 to L.R.C, and F31 AI194603 to L.R.C.). E.V. is a Howard Hughes Medical Institute Investigator. This study includes X-ray crystallography data collected at the Stanford Synchrotron Radiation Lightsource, SLAC National Accelerator Laboratory, which is supported by the U.S. Department of Energy, Office of Science, Office of Basic Energy Sciences under Contract No. DE-AC02-76SF00515. The SSRL Structural Molecular Biology Program is supported by the DOE Office of Biological and Environmental Research, and by the National Institutes of Health, National Institute of General Medical Sciences (P30GM133894). The contents of this publication are solely the responsibility of the authors and do not necessarily represent the official views of NIGMS or NIH. This study also includes X-ray crystallography data collected at the Advanced Photon Source, a U.S. Department of Energy (DOE) Office of Science User Facility operated for the DOE Office of Science by Argonne National Laboratory under Contract No. DE-AC02-06CH11357. Data were collected on the Northeastern Collaborative Access Team beamlines, which are funded by the National Institute of General Medical Sciences from the National Institutes of Health (P30 GM124165).

## Author Contributions

L.R.C. performed biochemical and structural experiments, generated figures, and wrote the manuscript. P.R. performed ChIP-Seq experiments. R.K.M. performed bacteriophage protection experiments with VapKSC and Bas42. E.V. obtained funding, consulted on experimental design, and wrote the manuscript. K.D.C. obtained funding, oversaw all experiments, assisted with structure determination, generated figures, and wrote the manuscript.

## Declaration of Interests

The authors declare no competing interests.

## Supplemental Information

**Table S1. Transcriptional regulators associated with Bub operons**

*(see separate Excel workbook)*

**Table S2. Identified CapK and CapS proteins**

*(see separate Excel workbook)*

**Table S3. Identified VapK, VapS, and VapC proteins**

*(see separate Excel workbook)*

**Table S4.**
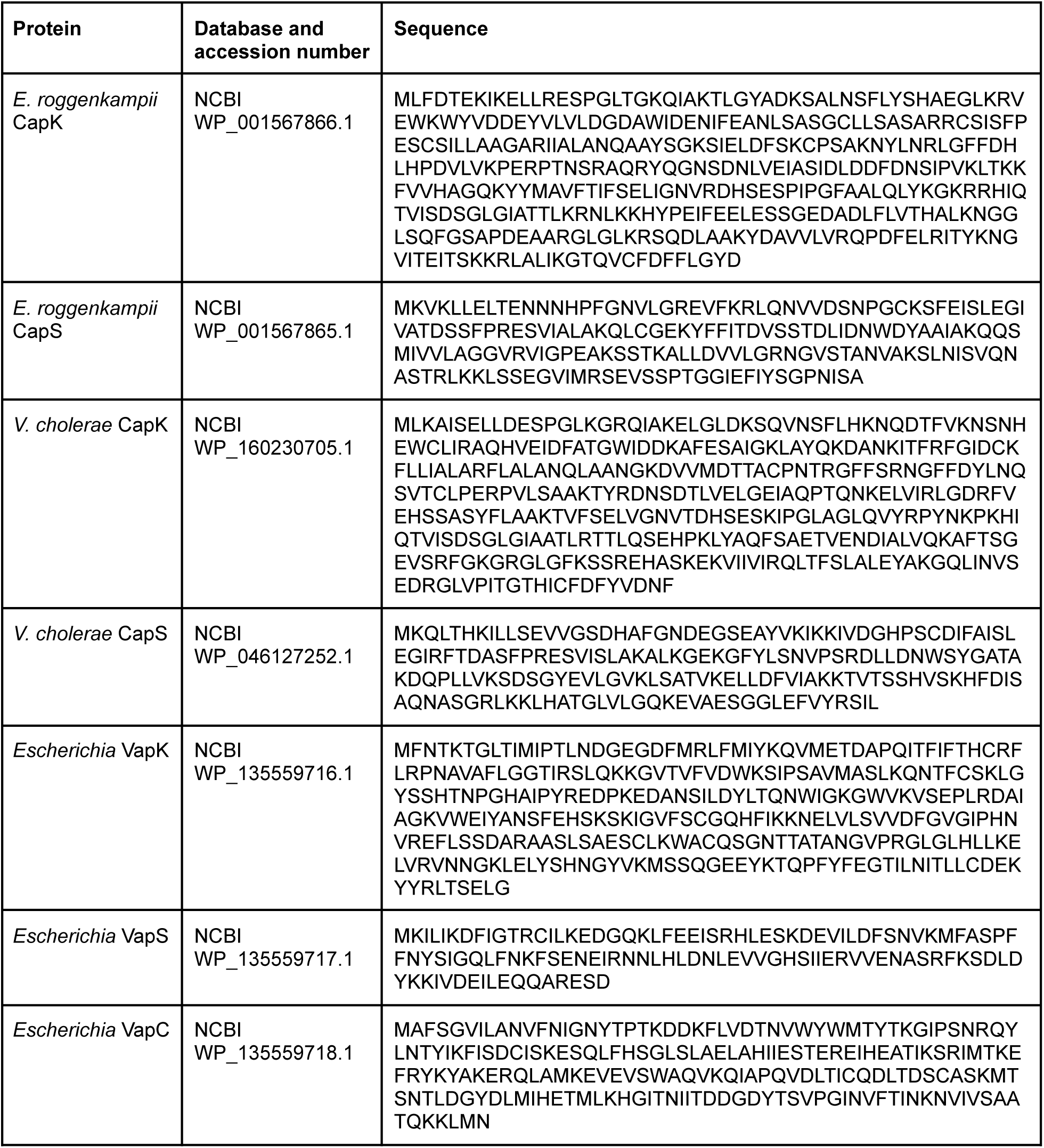
Proteins used in this study.

**Table S5.**
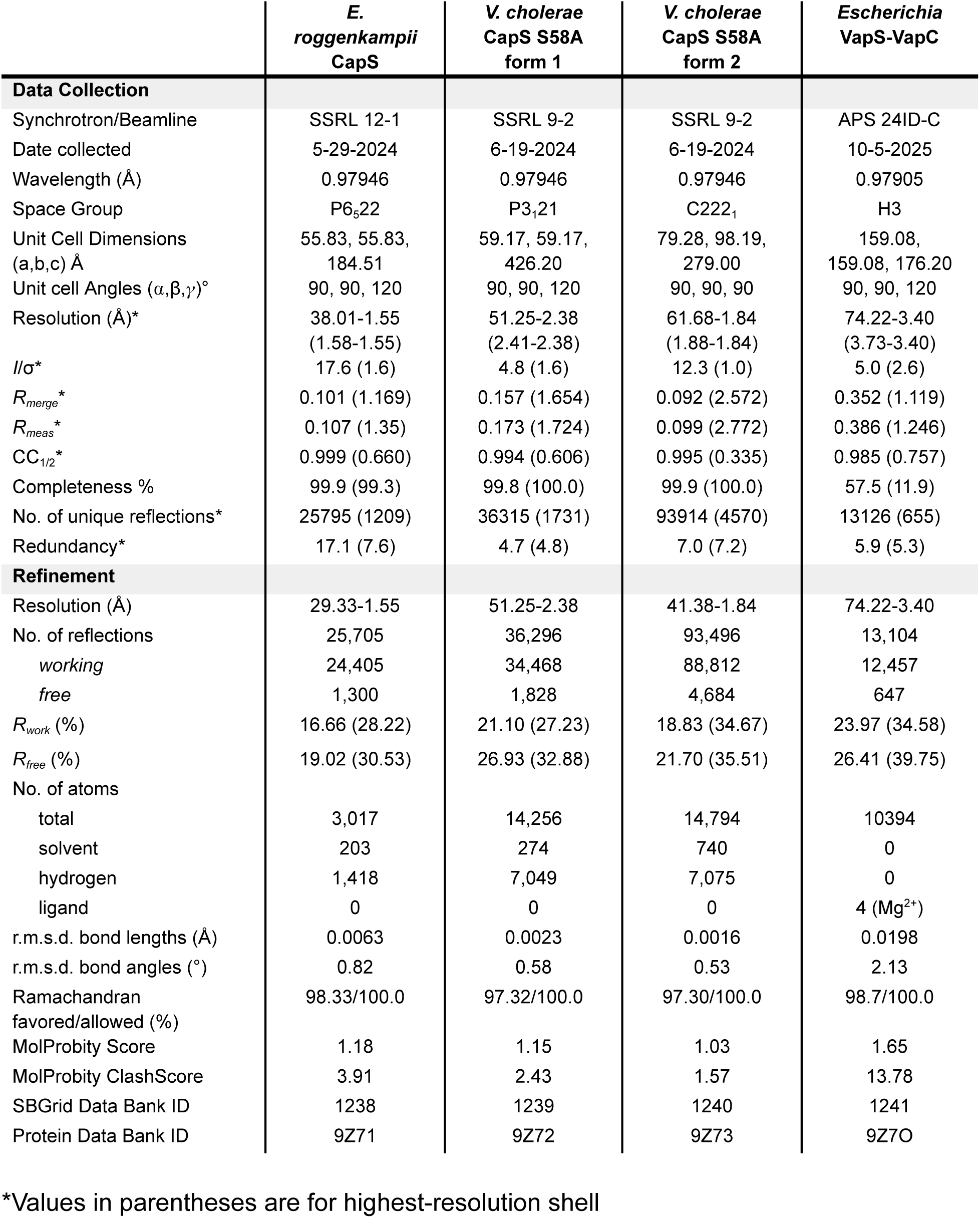
Crystallographic data collection and refinement statistics.

## SUPPLEMENTAL FIGURES

**Figure S1.**
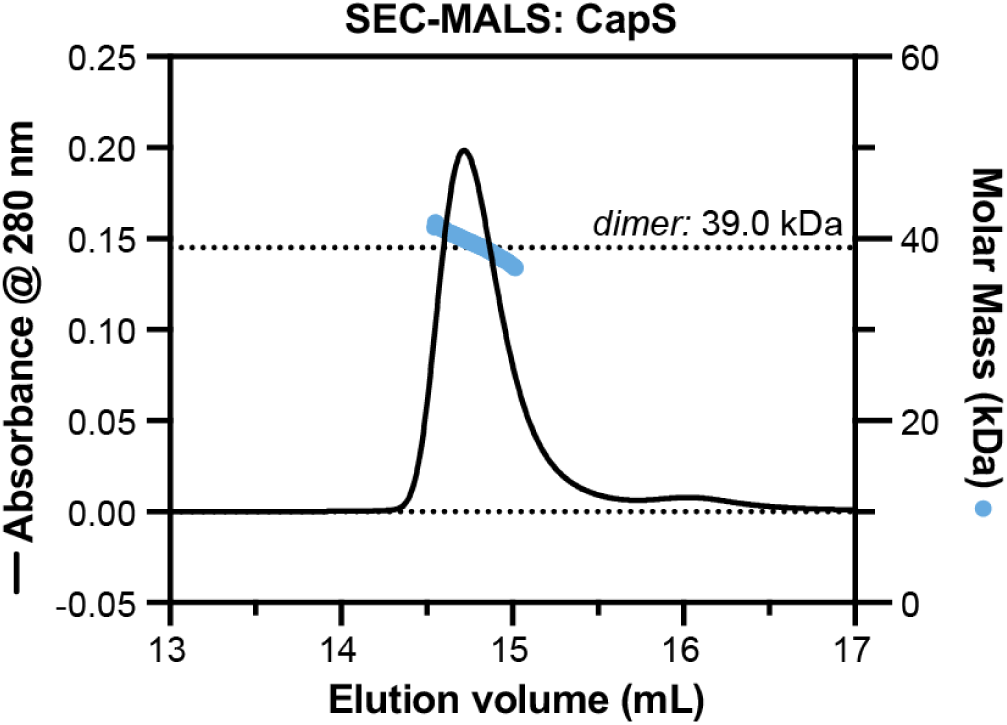
CapS SEC-MALS. Size exclusion chromatography coupled to multi-angle light scattering (SEC-MALS) analysis of purified *E. roggenkampii* CapS.

**Figure S2.**
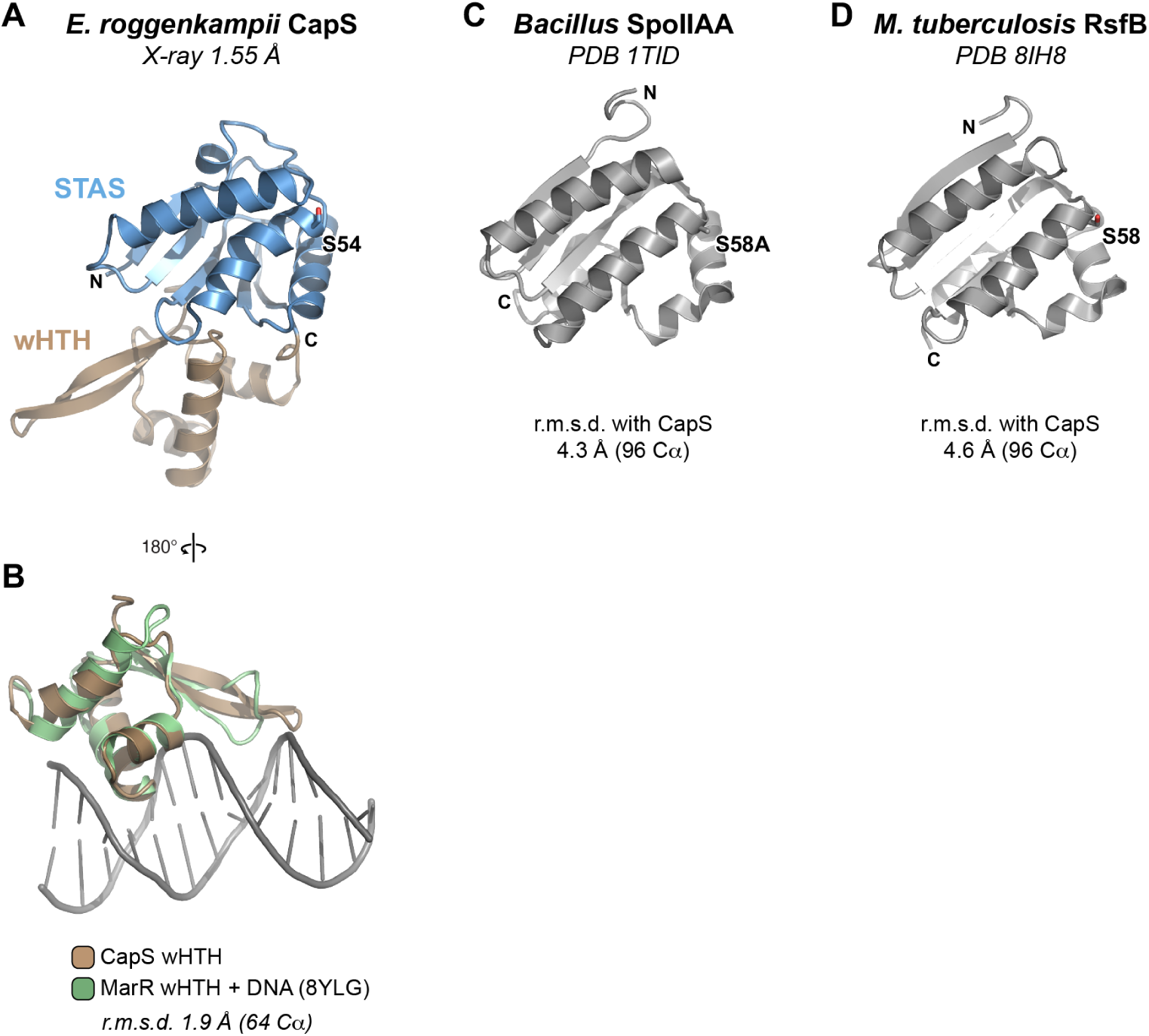
Structure of *E. roggenkampii* CapS. (A) Structure of *E. roggenkampii* CapS, with N-terminal STAS domain blue and C-terminal wHTH domain brown. The N- and C-termini of the STAS domain are labeled, and Ser54 is shown as sticks and labeled. (B) Overlay of *E. roggenkampii* CapS (brown) with a crystal structure of MarR bound to DNA (green; PDB ID 8YLG) (Song *et al*, 2024); only the wHTH domains of both proteins are shown. (C) Structure of *Geobacillus stearothermophilus* SpoIIAA (from PDB ID 1TID) (Masuda *et al*, 2004), with Ser58 (mutated to alanine) shown as sticks and labeled. The overall Cα r.m.s.d. (root mean squared deviation) with CapS is 4.3 Å over 96 residues. (D) Structure of *Mycobacterium tuberculosis* RsfB (from PDB ID 8IH8; unpublished), with Ser58 shown as sticks and labeled. The overall Cα r.m.s.d. with CapS is 4.6 Å over 96 residues.

**Figure S3.**
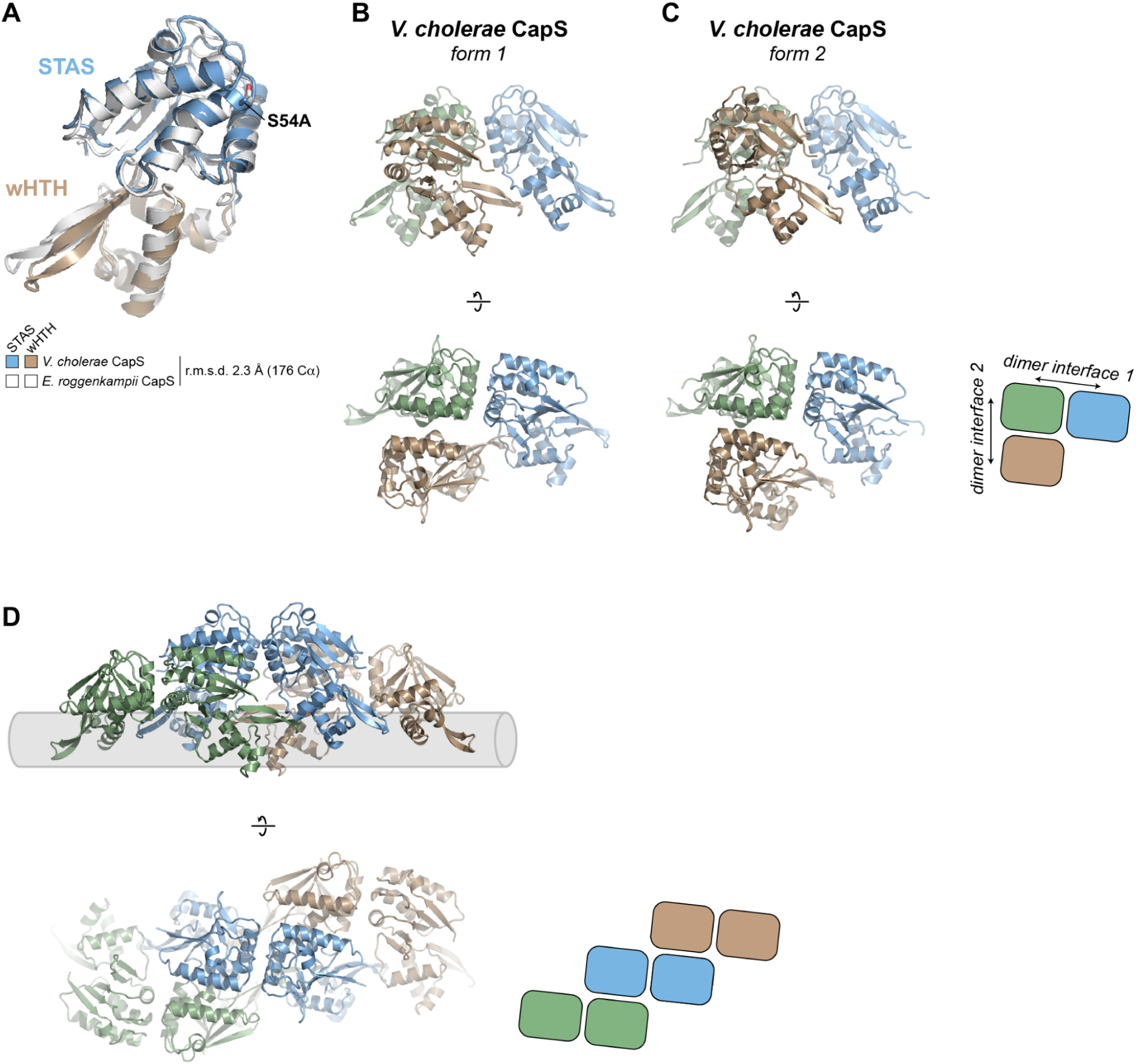
Structure of *V. cholerae* CapS (S58A) (A) Overlay of *V. cholerae* CapS (STAS domain blue; wHTH domain brown) from crystal form 1 with *E. roggenkampii* CapS (Cα r.m.s.d. 2.3 Å over 176 residues). (B) Two views of three non-crystallographic symmetry-related *V. cholerae* CapS protomers in crystal form 1, showing dimer interface 1 (green-blue) and dimer interface 2 (green-brown). (C) Two views of three non-crystallographic symmetry-related *V. cholerae* CapS protomers in crystal form 2, showing dimer interface 1 (green-blue) and dimer interface 2 (green-brown). (D) Two views of a *V. cholerae* CapS filament (six protomers) assembled from a combination of crystallographic and non-crystallographic symmetry. Three dimers assembled via interface 1 are colored green, blue, and brown, respectively. Each of these dimers associates with other dimer view interface 2. Shown in gray is a theoretical DNA duplex positioned such that all six CapS wHTH domains could bind it.

**Figure S4.**
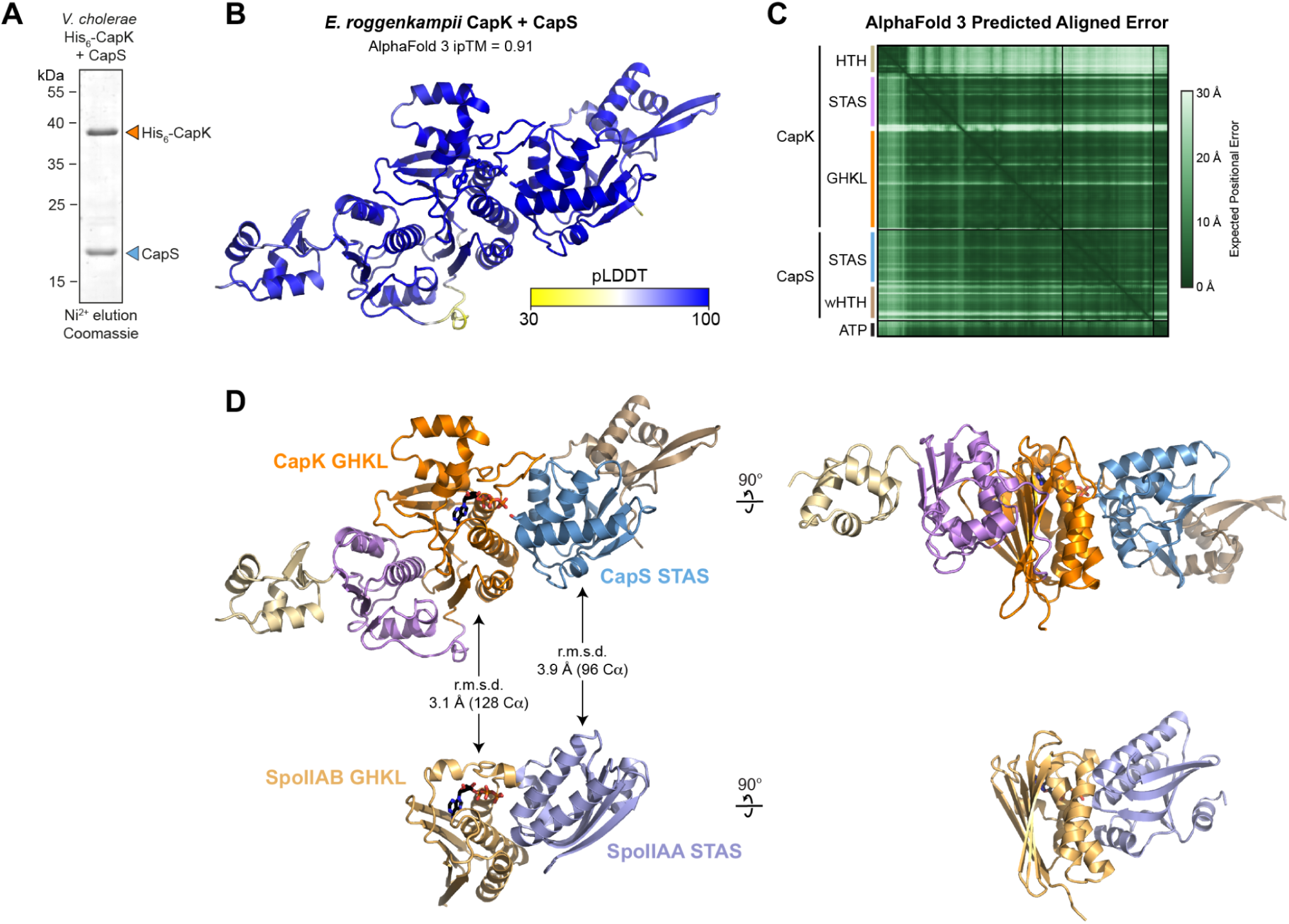
CapK and CapS resemble bacterial anti-sigma factors and their antagonists. (A) Ni^2+^ elution of co-expressed *V. cholerae* His_6_-CapK and untagged CapS. (B) AlphaFold 3 predicted structure of *E. roggenkampii* CapK bound to CapS, colored by confidence (pLDDT). (C) Predicted Aligned Error (PAE) plot for the model shown in panel (A). (D) *Top:* Two views of the AlphaFold 3 predicted structure of *E. roggenkampii* CapK bound to CapS, colored by domain as in **Figure 4A**. ATP bound to the CapK GHKL domain is shown in sticks. *Bottom:* Two views of the *Bacillus* SpoIIAB-SpoIIAA complex (PDB ID 1TID) (Masuda *et al*, 2004), aligned to CapK-CapS. ATP bound to the SpoIIAB GHKL domain is shown in sticks.

**Figure S5.**
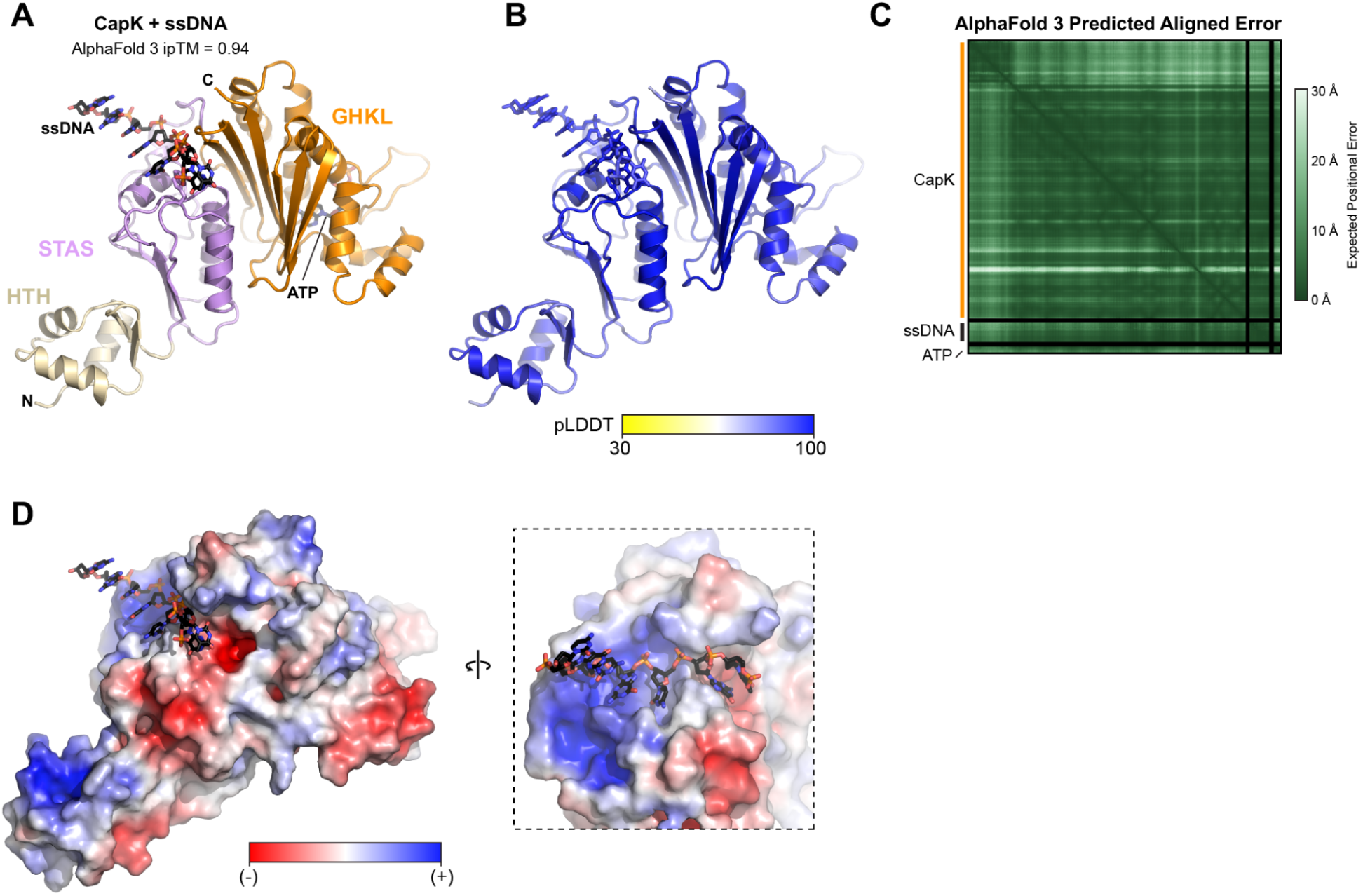
AlphaFold 3 predicted model of the *E. roggenkampii* CapK-ssDNA complex. (A) AlphaFold 3 predicted model of a complex of *E. roggenkampii* CapK (HTH domain yellow, STAS domain pink, GHKL domain orange), poly-T ssDNA (black), ATP (black), and Mg^2+^ (gray; not visible). CapK Trp68 is shown as sticks. (B) View as in panel (A), colored by confidence (pLDDT). (C) Predicted Aligned Error (PAE) plot for the model shown in panel (A). (D) Two views the predicted CapK-ssDNA structure, with CapK shown as a molecular surface and colored by charge (negative charge red, positive charge blue).

**Figure S6.**
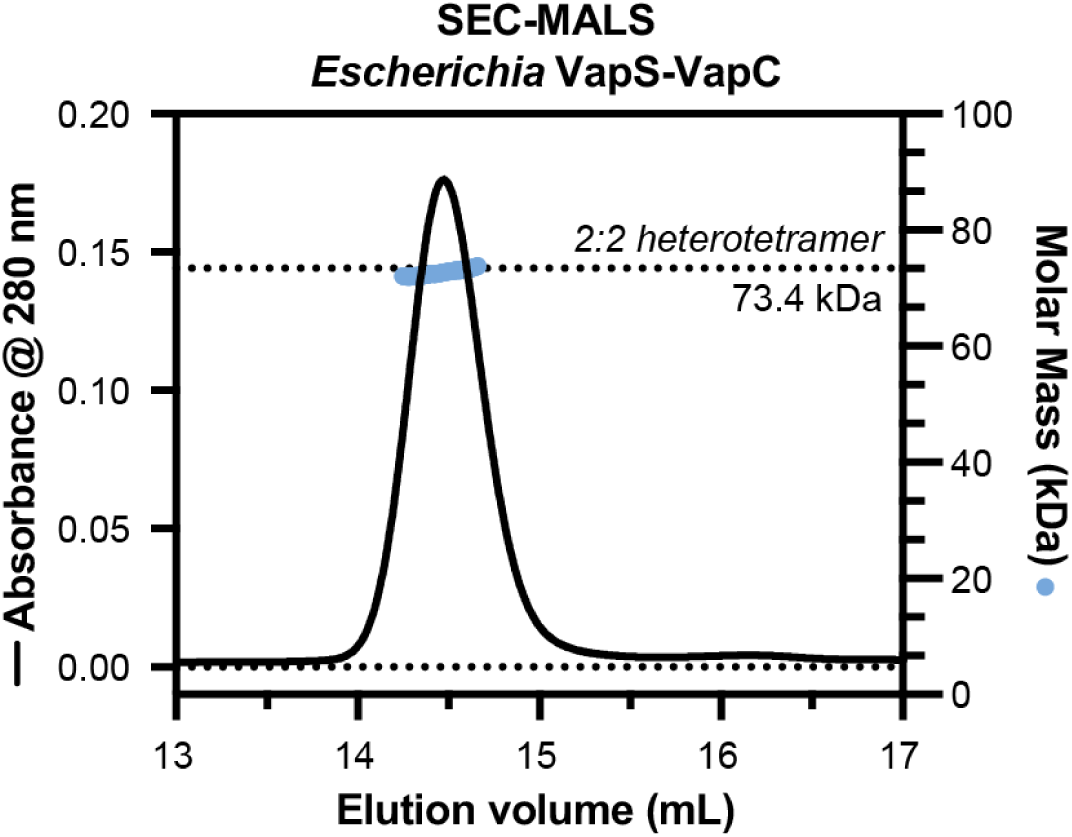
SEC-MALS of *Escherichia* VapS-VapC. Size exclusion chromatography coupled to multi-angle light scattering (SEC-MALS) of the Escherichia VapS-VapC complex. Black line indicates absorbance at 280 nm, and blue circles indicate measured molecular weight. A dotted line indicates the molecular weight of a 2:2 heterotetramer of VapS and VapC.

**Figure S7.**
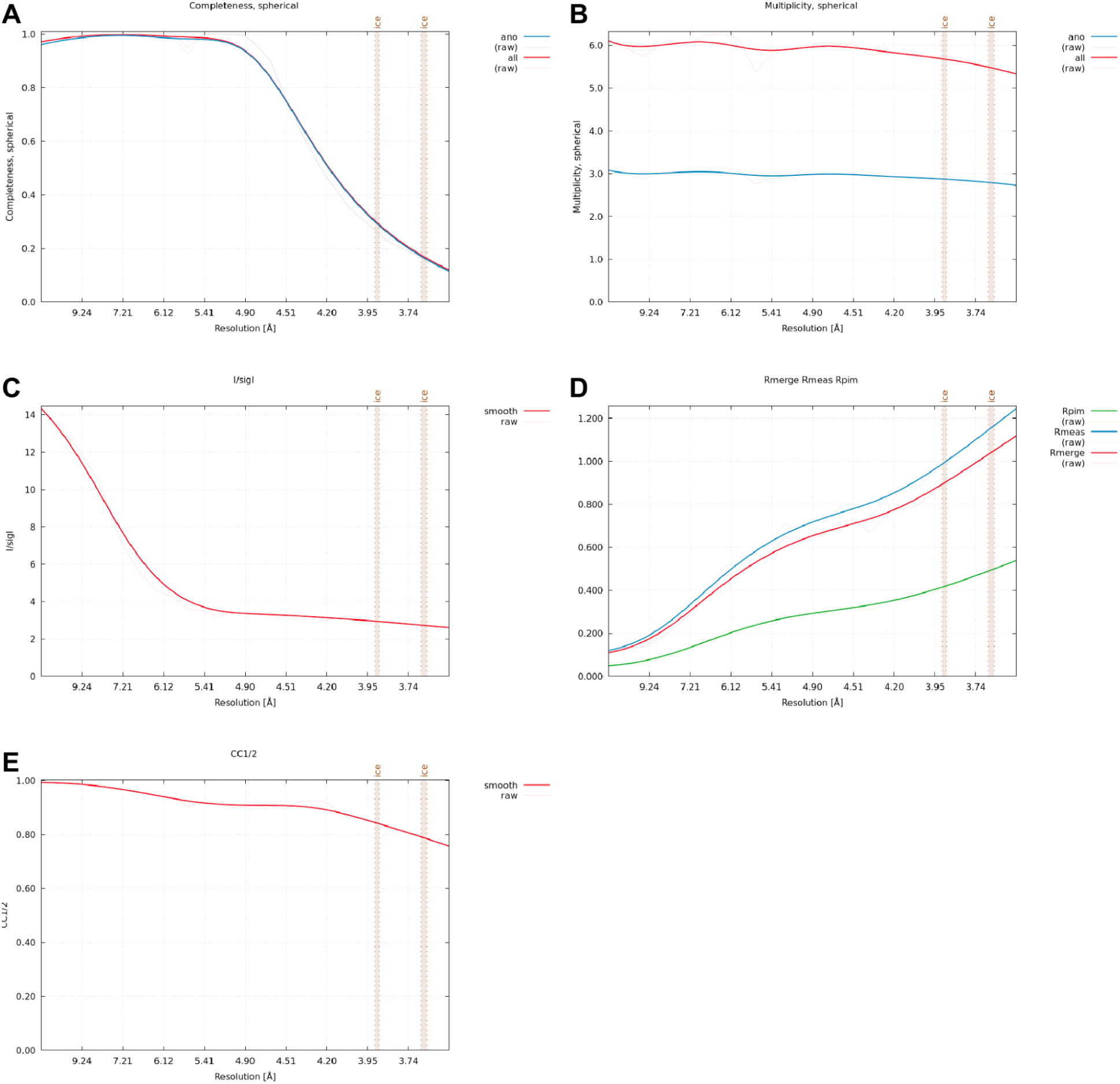
Anisotropic processing of *Escherichia* VapS-VapC crystallographic data. (A) Graph of completeness versus resolution for the STARANISO-processed X-ray diffraction dataset of *Escherichia* VapS-VapC. Red line indicates native completeness; blue line indicates anomalous completeness. Brown vertical bars indicate resolution ranges typical of ice crystal diffraction. (B) Graph of multiplicity versus resolution for the STARANISO-processed X-ray diffraction dataset of *Escherichia* VapS-VapC. (C) Graph of intensity (I/σI) versus resolution for the STARANISO-processed X-ray diffraction dataset of *Escherichia* VapS-VapC. (D) Graph of *R* values versus resolution for the STARANISO-processed X-ray diffraction dataset of *Escherichia* VapS-VapC. Red line indicates *R_merge_*; blue line indicates *R_meas_*, and green line indicates *R_pim_*. (E) Graph of CC_1/2_ versus resolution for the STARANISO-processed X-ray diffraction dataset of *Escherichia* VapS-VapC.

**Figure S8.**
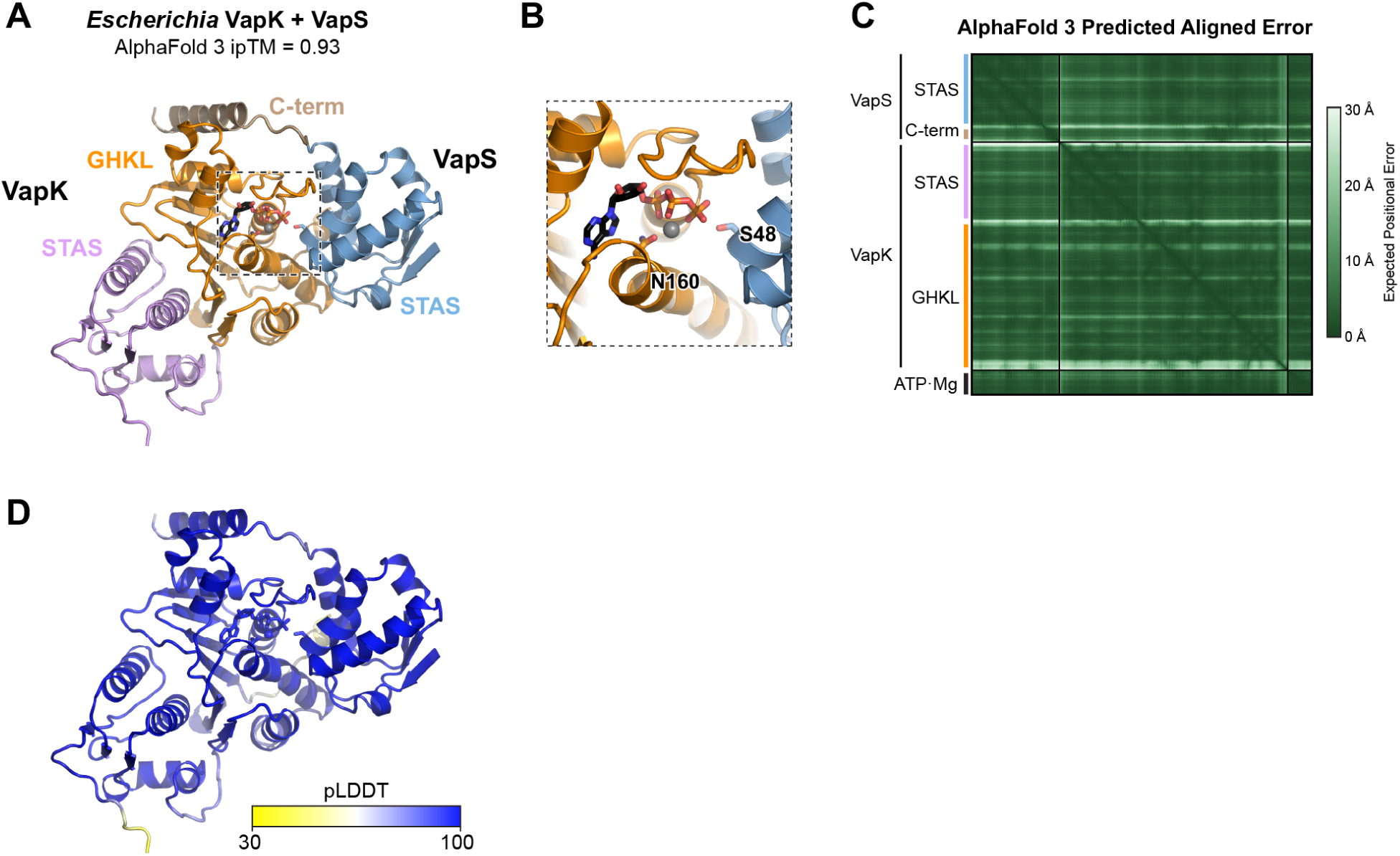
AlphaFold 3 predicted model of the *Escherichia* VapK-VapS complex. (A) AlphaFold 3 predicted model of a complex of *Escherichia* VapK (STAS domain pink, GHKL domain orange), VapS (STAS domain blue, C-terminal region brown), ATP (black), and Mg^2+^ (gray). (B) Closeup of the VapK GHKL kinase active site, with VapK Asn160 and VapS Ser48 shown as sticks. (C) AlphaFold 3 predicted aligned error (PAE) plot for the prediction shown in panel (A). (D) View as in panel (A), colored by confidence (pLDDT).

**Figure S9.**
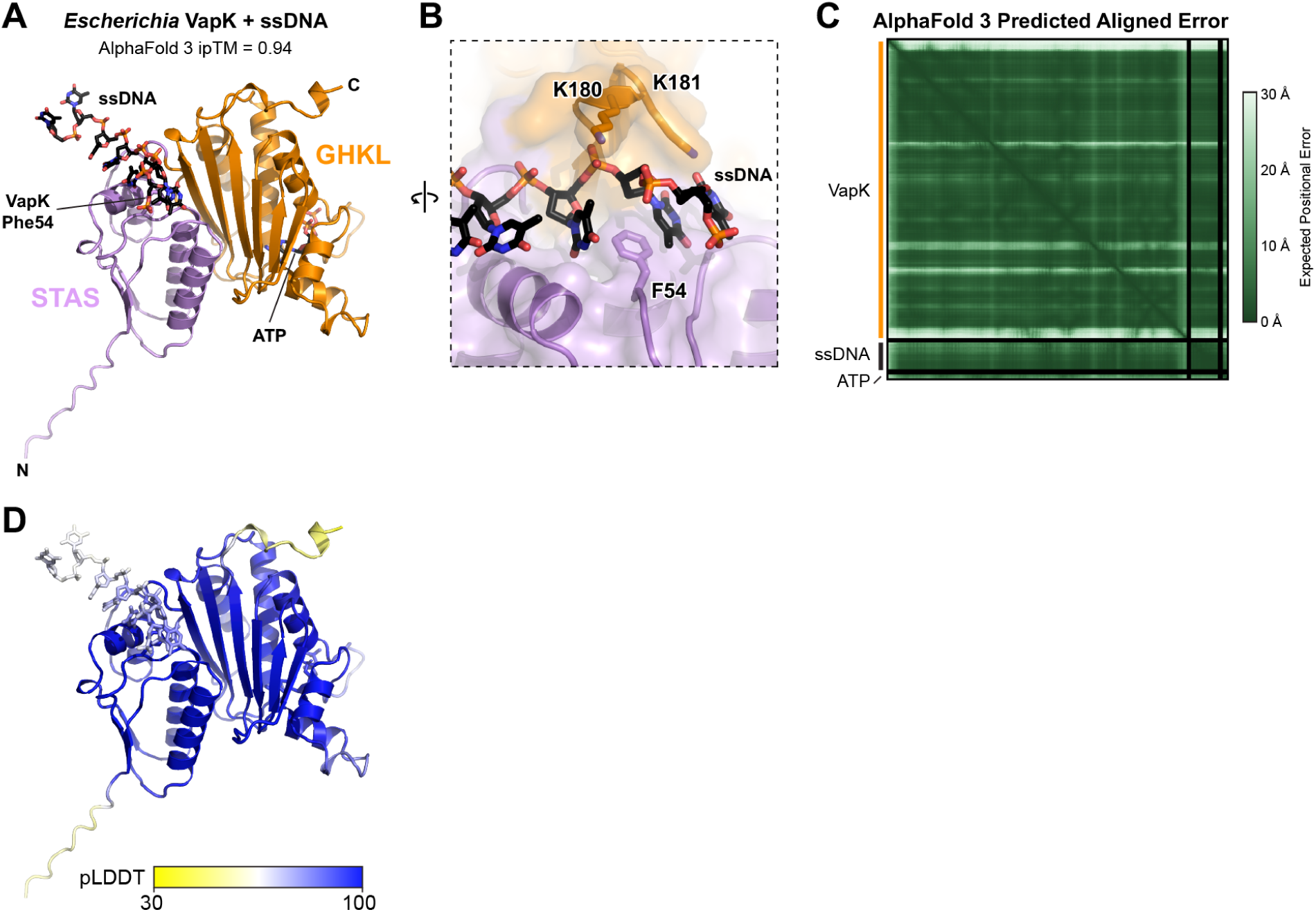
AlphaFold 3 predicted model of the *Escherichia* VapK-ssDNA complex. (A) AlphaFold 3 predicted model of a complex of *Escherichia* VapK (STAS domain pink, GHKL domain orange), a 7mer poly-T ssDNA (black), ATP (black), and Mg^2+^ (gray; not visible). VapK Phe54 is shown as sticks. (B) Closeup of ssDNA binding by VapK, with Phe54, Lys180, and Lys181 shown as sticks and labeled. (C) Predicted Aligned Error (PAE) plot for the model shown in panel (A). (D) View as in panel (A), colored by confidence (pLDDT).

## Notes

### Competing Interest Statement

The authors have declared no competing interest.

### Summary of Updates

Updating typo in first author name, adding funding information, and adding supplemental tables 1-3 Excel file.

## References

Adams PD, Afonine PV, Bunkóczi G, Chen VB, Davis IW, Echols N, Headd JJ, Hung L-W, Kapral GJ, Grosse-Kunstleve RW, et al (2010) PHENIX: a comprehensive Python-based system for macromolecular structure solution. Acta Crystallogr D Biol Crystallogr 66: 213–221

Amann E, Ochs B & Abel KJ (1988) Tightly regulated tac promoter vectors useful for the expression of unfused and fused proteins in Escherichia coli. Gene 69: 301–315

Aravind L & Koonin EV (2000) The STAS domain - a link between anion transporters and antisigma-factor antagonists. Curr Biol 10: R53–5

Arcus VL, McKenzie JL, Robson J & Cook GM (2011) The PIN-domain ribonucleases and the prokaryotic VapBC toxin-antitoxin array. Protein Eng Des Sel 24: 33–40

Baba T, Ara T, Hasegawa M, Takai Y, Okumura Y, Baba M, Datsenko KA, Tomita M, Wanner BL & Mori H (2006) Construction of Escherichia coli K-12 in-frame, single-gene knockout mutants: the Keio collection. Mol Syst Biol 2: 2006.0008

Bernheim A & Sorek R (2020) The pan-immune system of bacteria: antiviral defence as a community resource. Nat Rev Microbiol 18: 113–119

Blankenchip CL & Corbett KD (2024) Bacterial WYL domain transcriptional repressors sense single-stranded DNA to control gene expression. Nucleic Acids Res 52: 13723–13732

Blankenchip CL, Nguyen JV, Lau RK, Ye Q, Gu Y & Corbett KD (2022) Control of bacterial immune signaling by a WYL domain transcription factor. Nucleic Acids Res 50: 5239–5250

Chambers LR, Ye Q, Cai J, Gong M, Ledvina HE, Zhou H, Whiteley AT, Suhandynata RT & Corbett KD (2024) A eukaryotic-like ubiquitination system in bacterial antiviral defence. Nature 631: 843–849

Cohen D, Melamed S, Millman A, Shulman G, Oppenheimer-Shaanan Y, Kacen A, Doron S, Amitai G & Sorek R (2019) Cyclic GMP-AMP signalling protects bacteria against viral infection. Nature 574: 691–695

Conway JM, Walton WG, Salas-González I, Law TF, Lindberg CA, Crook LE, Kosina SM, Fitzpatrick CR, Lietzan AD, Northen TR, et al (2022) Diverse MarR bacterial regulators of auxin catabolism in the plant microbiome. Nat Microbiol 7: 1817–1833

Danecek P, Bonfield JK, Liddle J, Marshall J, Ohan V, Pollard MO, Whitwham A, Keane T, McCarthy SA, Davies RM, et al (2021) Twelve years of SAMtools and BCFtools. Gigascience 10

Datsenko KA & Wanner BL (2000) One-step inactivation of chromosomal genes in Escherichia coli K-12 using PCR products. Proc Natl Acad Sci U S A 97: 6640–6645

Davies BW, Bogard RW, Young TS & Mekalanos JJ (2012) Coordinated regulation of accessory genetic elements produces cyclic di-nucleotides for V. cholerae virulence. Cell 149: 358–370

Dienemann C, Bøggild A, Winther KS, Gerdes K & Brodersen DE (2011) Crystal structure of the VapBC toxin-antitoxin complex from Shigella flexneri reveals a hetero-octameric DNA-binding assembly. J Mol Biol 414: 713–722

Dutta R & Inouye M (2000) GHKL, an emergent ATPase/kinase superfamily. Trends Biochem Sci 25: 24–28

Dworkin J & Losick R (2001) Differential gene expression governed by chromosomal spatial asymmetry. Cell 107: 339–346

Emsley P, Lohkamp B, Scott WG & Cowtan K (2010) Features and development of Coot. Acta Crystallogr D Biol Crystallogr 66: 486–501

Evans PR & Murshudov GN (2013) How good are my data and what is the resolution? Acta Crystallogr D Biol Crystallogr 69: 1204–1214

Gao Y, Cao D, Zhu J, Feng H, Luo X, Liu S, Yan X-X, Zhang X & Gao P (2020) Structural insights into assembly, operation and inhibition of a type I restriction-modification system. Nat Microbiol 5: 1107–1118

Georjon H & Bernheim A (2023) The highly diverse antiphage defence systems of bacteria. Nat Rev Microbiol 21: 686–700

Gong M, Ye Q, Gu Y, Chambers LR, Bobkov AA, Arakawa NK, Matyszewski M & Corbett KD (2025) Structural diversity and oligomerization of bacterial ubiquitin-like proteins. Structure

Hampton HG, Watson BNJ & Fineran PC (2020) The arms race between bacteria and their phage foes. Nature 577: 327–336

Hör J, Wolf SG & Sorek R (2024) Bacteria conjugate ubiquitin-like proteins to interfere with phage assembly. Nature 631: 850–856

Høyland-Kroghsbo NM, Paczkowski J, Mukherjee S, Broniewski J, Westra E, Bondy-Denomy J & Bassler BL (2017) Quorum sensing controls the Pseudomonas aeruginosa CRISPR-Cas adaptive immune system. Proc Natl Acad Sci U S A 114: 131–135

Johnson AG, Wein T, Mayer ML, Duncan-Lowey B, Yirmiya E, Oppenheimer-Shaanan Y, Amitai G, Sorek R & Kranzusch PJ (2022) Bacterial gasdermins reveal an ancient mechanism of cell death. Science 375: 221–225

Kabsch W (2010) XDS. Acta Crystallogr D Biol Crystallogr 66: 125–132

Kang S-M, Jin C, Kim D-H, Lee Y & Lee B-J (2020) Structural and functional study of the Klebsiella pneumoniae VapBC toxin-antitoxin system, including the development of an inhibitor that activates VapC. J Med Chem 63: 13669–13679

Kim A-R, Hu Y, Comjean A, Rodiger J, Mohr SE & Perrimon N (2024) Enhanced Protein-Protein Interaction Discovery via AlphaFold-Multimer. bioRxiv: 2024.02.19.580970

Krissinel E & Henrick K (2007) Inference of macromolecular assemblies from crystalline state. J Mol Biol 372: 774–797

Langmead B & Salzberg SL (2012) Fast gapped-read alignment with Bowtie 2. Nat Methods 9: 357–359

Langmead B, Trapnell C, Pop M & Salzberg SL (2009) Ultrafast and memory-efficient alignment of short DNA sequences to the human genome. Genome Biol 10: R25

Lau RK, Enustun E, Gu Y, Nguyen JV & Corbett KD (2022) A conserved signaling pathway activates bacterial CBASS immune signaling in response to DNA damage. EMBO J 41: e111540

Li H, Handsaker B, Wysoker A, Fennell T, Ruan J, Homer N, Marth G, Abecasis G, Durbin R & 1000 Genome Project Data Processing Subgroup (2009) The Sequence Alignment/Map format and SAMtools. Bioinformatics 25: 2078–2079

Lopatina A, Tal N & Sorek R (2020) Abortive infection: Bacterial suicide as an antiviral immune strategy. Annu Rev Virol 7: 371–384

Luyten YA, Hausman DE, Young JC, Doyle LA, Higashi KM, Ubilla-Rodriguez NC, Lambert AR, Arroyo CS, Forsberg KJ, Morgan RD, et al (2022) Identification and characterization of the WYL BrxR protein and its gene as separable regulatory elements of a BREX phage restriction system. Nucleic Acids Res 50: 5171–5190

Maffei E, Shaidullina A, Burkolter M, Heyer Y, Estermann F, Druelle V, Sauer P, Willi L, Michaelis S, Hilbi H, et al (2021) Systematic exploration of Escherichia coli phage-host interactions with the BASEL phage collection. PLoS Biol 19: e3001424

Mariano G & Blower TR (2023) Conserved domains can be found across distinct phage defence systems. Mol Microbiol 120: 45–53

Masuda S, Murakami KS, Wang S, Anders Olson C, Donigian J, Leon F, Darst SA & Campbell EA (2004) Crystal structures of the ADP and ATP bound forms of the Bacillus anti-sigma factor SpoIIAB in complex with the anti-anti-sigma SpoIIAA. J Mol Biol 340: 941–956

McCoy AJ, Grosse-Kunstleve RW, Adams PD, Winn MD, Storoni LC & Read RJ (2007) Phaser crystallographic software. J Appl Crystallogr 40: 658–674

Miallau L, Faller M, Chiang J, Arbing M, Guo F, Cascio D & Eisenberg D (2009) Structure and proposed activity of a member of the VapBC family of toxin-antitoxin systems. VapBC-5 from Mycobacterium tuberculosis: VapBC-5 FROM MYCOBACTERIUM TUBERCULOSIS. J Biol Chem 284: 276–283

Millman A, Melamed S, Leavitt A, Doron S, Bernheim A, Hör J, Garb J, Bechon N, Brandis A, Lopatina A, et al (2022) An expanded arsenal of immune systems that protect bacteria from phages. Cell Host Microbe 30: 1556–1569.e5

Min AB, Miallau L, Sawaya MR, Habel J, Cascio D & Eisenberg D (2012) The crystal structure of the Rv0301-Rv0300 VapBC-3 toxin-antitoxin complex from M. tuberculosis reveals a Mg^2+^ ion in the active site and a putative RNA-binding site: Crystal Structure of the Rv0301-Rv0300 Complex. Protein Sci 21: 1754–1767

Montgomery MT, Guerrero Bustamante CA, Dedrick RM, Jacobs-Sera D & Hatfull GF (2019) Yet more evidence of collusion: A new viral defense system encoded by Gordonia phage CarolAnn. MBio 10

Moy BE & Seshu J (2021) STAS domain only proteins in bacterial gene regulation. Front Cell Infect Microbiol 11: 679982

Negri A, Werbowy O, Wons E, Dersch S, Hinrichs R, Graumann PL & Mruk I (2021) Regulator-dependent temporal dynamics of a restriction-modification system’s gene expression upon entering new host cells: single-cell and population studies. Nucleic Acids Res 49: 3826–3840

Oshiro RT, Dunham DT, Gill C, Chouldjian A, Piya D, Mutalik VK & Seed KD (2025) Surviving phage attack dynamically regulates bacterial immunity to defeat counterdefenses. bioRxiv

Pandey DP & Gerdes K (2005) Toxin-antitoxin loci are highly abundant in free-living but lost from host-associated prokaryotes. Nucleic Acids Res 33: 966–976

Parma DH, Snyder M, Sobolevski S, Nawroz M, Brody E & Gold L (1992) The Rex system of bacteriophage lambda: tolerance and altruistic cell death. Genes Dev 6: 497–510

Patterson AG, Jackson SA, Taylor C, Evans GB, Salmond GPC, Przybilski R, Staals RHJ & Fineran PC (2016) Quorum sensing controls adaptive immunity through the regulation of multiple CRISPR-Cas systems. Mol Cell 64: 1102–1108

Payne LJ, Meaden S, Mestre MR, Palmer C, Toro N, Fineran PC & Jackson SA (2022) PADLOC: a web server for the identification of antiviral defence systems in microbial genomes. Nucleic Acids Res 50: W541–W550

Peng JW, Yuan H & Tan XS (2017) Crystal structure of the multiple antibiotic resistance regulator MarR from Clostridium difficile. Acta Crystallogr F Struct Biol Commun 73: 363–368

Picton DM, Harling-Lee JD, Duffner SJ, Went SC, Morgan RD, Hinton JCD & Blower TR (2022) A widespread family of WYL-domain transcriptional regulators co-localizes with diverse phage defence systems and islands. Nucleic Acids Res 50: 5191–5207

Prabaharan C, Kandavelu P, Packianathan C, Rosen BP & Thiyagarajan S (2019) Structures of two ArsR As(III)-responsive transcriptional repressors: Implications for the mechanism of derepression. J Struct Biol 207: 209–217

Price MN & Arkin AP (2024) A fast comparative genome browser for diverse bacteria and archaea. PLoS One 19: e0301871

Rousset F, Osterman I, Scherf T, Falkovich AH, Leavitt A, Amitai G, Shir S, Malitsky S, Itkin M, Savidor A, et al (2025) TIR signaling activates caspase-like immunity in bacteria. Science 387: 510–516

Saha CK, Sanches Pires R, Brolin H, Delannoy M & Atkinson GC (2021) FlaGs and webFlaGs: discovering novel biology through the analysis of gene neighbourhood conservation. Bioinformatics 37: 1312–1314

Solovyev A & Salamov A (2011) Automatic Annotation of Microbial Genomes and Metagenomic Sequences. In Metagenomics and its Applications in Agriculture, Biomedicine and Environmental Studies, Li RW (ed) pp 61–78. Hauppauge, NY: Nova Biomedical

Song WS, Ki DU, Cho HY, Kwon OH, Cho H & Yoon S-I (2024) Structural basis of transcriptional regulation by UrtR in response to uric acid. Nucleic Acids Res 52: 13192–13205

Tamulaitiene G, Sabonis D, Sasnauskas G, Ruksenaite A, Silanskas A, Avraham C, Ofir G, Sorek R, Zaremba M & Siksnys V (2024) Activation of Thoeris antiviral system via SIR2 effector filament assembly. Nature 627: 431–436

Tomasz M (1994) The mitomycins: Natural cross-linkers of DNA. In Molecular Aspects of Anticancer Drug-DNA Interactions pp 312–349. London: Macmillan Education UK

Truglio JJ, Croteau DL, Van Houten B & Kisker C (2006) Prokaryotic nucleotide excision repair: the UvrABC system. Chem Rev 106: 233–252

Viswanathan T, Chen J, Wu M, An L, Kandavelu P, Sankaran B, Radhakrishnan M, Li M & Rosen BP (2021) Functional and structural characterization of AntR, an Sb(III) responsive transcriptional repressor. Mol Microbiol 116: 427–437

Ye Q, Gong M, Cai J, Chambers LR, Zhou H, Suhandynata RT & Corbett KD (2025) Mechanistic basis for protein conjugation in a diverged bacterial ubiquitination pathway. Nat Struct Mol Biol

Ye Q, Lau RK, Mathews IT, Birkholz EA, Watrous JD, Azimi CS, Pogliano J, Jain M & Corbett KD (2020) HORMA domain proteins and a Trip13-like ATPase regulate bacterial cGAS-like enzymes to mediate bacteriophage immunity. Mol Cell 77: 709–722.e7

